# Dual Recognition Drives Site-Directed G-Quadruplex Stabilization: Exploring Oligonucleotide Design in G4 Ligand-Oligonucleotide Conjugates

**DOI:** 10.64898/2026.04.08.717194

**Authors:** Alva Abrahamsson, Sakina Khwaja, Steven Vertueux, Andreas Berner, Koit Aasumets, Namrata Chaudhari, Chandan Kumar, Lotte Stietz, Tom Baladi, Anders Dahlén, Sjoerd Wanrooij, Erik Chorell

## Abstract

G-quadruplex (G4) DNA structures are increasingly recognized for their roles in key cellular processes, including transcriptional regulation and genome stability, making them attractive therapeutic targets. Selective recognition of individual G4s remains challenging due to the high structural similarity among human G4 motifs. The G4 Ligand-conjugated Oligonucleotide strategy addresses this need by combining the G4-binding capabilities of small-molecule G4-ligands with the sequence specificity of an oligonucleotide complementary to the flanking region of the target G4. Here, we systematically explore how the oligonucleotide component governs G4 binding and stabilization by varying its length, backbone composition, and sequence complementarity. This revealed that efficient G4 recognition depends on a strong interdependence between hybridization and G4-ligand binding, such that both elements cooperatively reinforce complex stability and site specificity. Central mismatches disrupt this dual recognition and reduce selectivity. While longer oligonucleotides hybridize more slowly, they form more stable complexes and show stronger G4 stabilization in thermal melting and polymerase stop assays. Replacing the DNA oligonucleotide with peptide nucleic acid enhances binding strength, thermal stability, and metabolic stability, but selective G4 stabilization is achieved only upon ligand conjugation. Together, these results show how rational oligonucleotide design enables selective and potent recognition of G4 structures using GL-Os.

## Introduction

G-quadruplex (G4) DNA structures are non-canonical secondary conformations that form either intra- or intermolecularly within guanine-rich regions of the genome. These structures are characterized by the formation of G-tetrads, which are planar arrangements of four guanine bases stabilized by Hoogsteen hydrogen bonding. G-tetrads can self-stack through arene-arene interactions and are further stabilized by monovalent or divalent cations, such as Na^+^ or K^+^.^1^ G4 structures exhibit significant topological polymorphism, with their conformation influenced by factors such as strand orientation, loop length and the nature of the stabilizing cation.^2^

G4 formation within promotor regions has been greatly investigated through *in vitro* and cellular assays, where these structures have demonstrated a direct role in transcriptional regulation.^3,4^ Among the most thoroughly studied examples is the oncogene *c-MYC*, which encodes the transcription factor MYC, and this G4 is of particular interest due to its potential as a therapeutic target in cancer.^5^ The guanine-rich nuclease hyper-sensitivity element III_I_, is responsible for the majority of the MYC transcription^5^, which contains a well-characterized G4-forming sequence termed Pu27. Small molecules, referred to as G4-ligands, have been shown to bind and stabilize this G4 structure, resulting in downregulation of the MYC expression.^6^ However, many G4-ligands act on multiple genomic sites, and the observed effects may reflect broader, genome-wide G4 stabilization rather than a promotor-specific response.^7,8^ This complexity is compounded by the potential formation of over 700.000 G4 structures throughout the human genome,^9^ with the G-tetrad as a conserved structural motif serving as the primary binding site for most G4-targeting ligands. As a result, there is a pressing need for strategies that enable selective targeting of individual G4 structures, to advance the developments of therapeutic agents and elucidating the cellular functions of G4s.

To achieve sequence-specific G4 recognition, one promising approach involves conjugating a G4-specific ligand to an oligonucleotide designed to be complementary to the flanking sequence of the target G4. This design guides the ligand to the intended G4 structure while minimizing off-target interactions with non-matching G4s.^10^ This approach is referred to as the G4-Ligand conjugated Oligonucleotide (GL-O) strategy (Figure 1A).^10^

**Figure 1.**
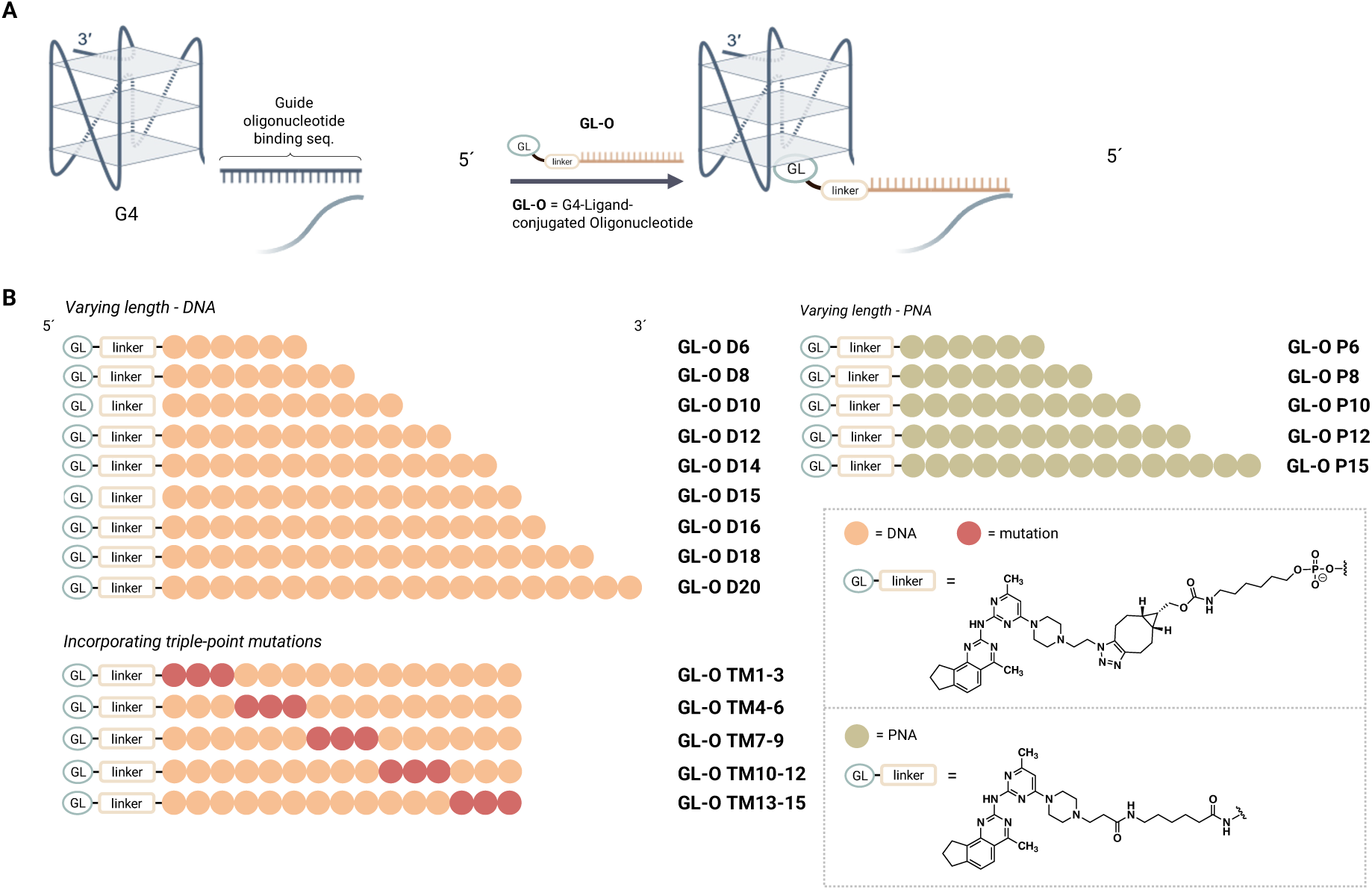
A) Overview of the G4 Ligand-conjugated Oligonucleotide (GL-Os) strategy and how the oligonucleotide binds to the 5’-flanking sequence of the G4. B) Modified GL-Os based on oligonucleotide length (GL-Os **D6**-**D20**), backbone composition (GL-Os **P6**-**P15**) and systematically introduced triple-point mutations (GL-Os **TM 1-3**, **TM4-6**, **TM7-9**, **TM10-12** and **TM13-15**).

Previous studies on the GL-O strategy have included detailed investigations into both the linker composition,^11^ and the G4-ligand,^12^ resulting in a flexible platform capable of accommodating various conjugation designs and fine-tuning the G4-ligand for optimal GL-O potency. Building on this foundation, the present study focuses on the primary determinant of the GL-O selectivity: the oligonucleotide. Specifically, we examine how three parameters; the oligonucleotide length, chemical backbone modifications and the degree of sequence complementarity, affect binding affinity, structural properties, and the ability to stabilize a target G4 DNA structure.

First, we investigated the effect of oligonucleotide length. The original GL-O design incorporated a 15-nucleotide (nt) oligonucleotide, selected based on foot printing experiments that identified a 14-15 nt single-stranded DNA region in the 5’-flanking sequence of the *c-MYC* promoter.^13^ However, a systematic evaluation of optimal oligonucleotide length has not yet been conducted. To address this, we preformed sequence alignments of various length oligonucleotides with three known G4 targets. This led us to design a series of GL-O constructs with oligonucleotide lengths ranging from 6 to 20-nt (Figure 1B).

Second, we evaluated how oligonucleotide composition influences GL-O performance. While selectivity requires sufficient length, this must be balanced against hybridization stability. To modulate these properties, we incorporated peptide nucleic acid (PNA), a DNA mimic with a neutral, peptide-like backbone developed by the Nielsen group in 1991.^14^ This uncharged backbone eliminates electrostatic repulsion during hybridization with the complementary DNA target, resulting in significantly enhanced binding affinity compared to unmodified DNA.^15^ In addition to their enhanced binding affinity, PNA oligonucleotides demonstrate pronounced metabolic stability against nuclease-mediated degradation.^15^ We therefore created a series of PNA-based GL-Os with oligonucleotide lengths ranging from 6 to 15-nt in length (Figure 1B).

Third, we assess the influence of sequence complementarity. A previous study examined the effects of central single-point mutations and double-point mutations in the peripheral regions.^10^ However, a more systematic analysis is still needed to identify clear trends in mismatch tolerance. To address this, we designed five GL-Os containing triple-point mutations at various positions along the oligonucleotide sequence (Figure 1B).

By systematically varying these three parameters, we aim to define the molecular principles governing G4 recognition and stabilization by GL-Os.

## Result

### Sequence conservation analysis of potential oligonucleotide lengths for various G4 targets

The length of the oligonucleotide is a key factor determining the selectivity of the GL-O strategy. A sufficiently long sequence enables the oligonucleotide to hybridize selectively to the desired genomic target while avoiding partial matches elsewhere. To evaluate how oligonucleotide length influences selectivity, we first aligned our target oligonucleotide binding sequences against the complete human genome. Genomic regions showing sequence similarity were then annotated to determine their genomic locations and whether they resided within coding or non-coding regions.

The analysis indicates that the required length of the oligonucleotide differs depending on the G4 target (Table 1). For the *c-MYC* Pu27 sequence, individual specificity could be retained with a 14-nt oligonucleotide, whereas the corresponding 13-nt variant showed complementarity to three additional genomic loci (listed in Table S4), none of which were adjacent to predicted G4-forming regions. In contrast, 13-nt sequences derived from *K-Ras* and *Helicase B* exhibited much lower selectivity, matching 74 and 309 genomic locations, respectively.

**Table 1.** Sequence conservation analysis of complementary sequences within the genome for three different G4 targets.

| Entry | Oligonucleotide length | Numbers of G4 flanking sequence sites in human genome |  |  |  |
| --- | --- | --- | --- | --- | --- |
|  |  | Random sequence | *c-Myc* Pu27 | *K-Ras* 32R | *Helicase B* G4-1 |
| 1 | 20 | No seq | 1 | 1 | 1 |
| 2 | 19 | No seq | 1 | 1 | 1 |
| 3 | 18 | No seq | 1 | 1 | 1 |
| 4 | 17 | No seq | 1 | 1 | 1 |
| 5 | 16 | No seq | 1 | 1 | 6 |
| 6 | 15 | 1 | 1 | 2 | 16 |
| 7 | 14 | 2 | 1 | 3 | 54 |
| 8 | 13 | 10 | 4 | 74 | 309 |
| 9 | 12 | 55 | 15 | 210 | 662 |
| 10 | 11 | 157 | 78 | 788 | 1838 |
| 11 | 10 | 927 | 330 | 2141 | 7228 |
| 12 | 9 | 5179 | 1816 | 8119 | 35586 |
| 13 | 8 | 21398 | 56285 | 22049 | 94195 |
| 14 | 7 | 85695 | 253973 | 59166 | 455120 |
| 15 | 6 | 390955 | 1556186 | 250643 | 1639320 |
| 16 | 5 | 1743835 | 5065989 | 1021355 | 5697262 |

### Design of Oligonucleotide modified GL-Os

The chemical design of the GL-O conjugate builds on previous work in which the G4-ligand and the linker chemistry were extensively investigated. In the present study, we retained the same G4-ligand as described in the initial report of the GL-O strategy.^10^ Conjugation was achieved through a strain-promoted azide-alkyne cycloaddition reaction, coupling an azide-functionalized G4-ligand with an oligonucleotide carrying a 5’-BCN.^11^ This approach provides a robust, copper-free biorthogonal linkage that ensure efficient conjugations.

For the DNA-based GL-Os, a series of oligonucleotides of varying lengths (6, 8, 10, 12, 14, 15, 16, 18 and 20-nt) were synthesized and conjugated to the G4-ligand to generate **GL-Os D6**-**D20** (see Table S2 for sequences and Table S3 for yields).

To evaluate the effect of backbone composition, a corresponding set of PNA-based GL-Os with different length oligonucleotides (6, 8, 10, 12, and 15-nt), named **GL-Os P6**-**P15**, were synthesized (Scheme 1). The PNA backbone oligonucleotide was synthesized on an automated solid-phase peptide synthesizer. PNA elongation was carried out in accordance with standard procedures.^16^ Conjugation of the G4-ligand to the *N*-terminus was performed directly on solid support under amide coupling conditions using the carboxylic acid analogue of the G4-ligand previously described (Scheme 1).^11^ Subsequent deprotection and cleavage using TFA/m-cresol yielded the purified **GL-Os P6-P15** (see Table S2 for sequences and Table S3 for yields).

**Scheme 1.**
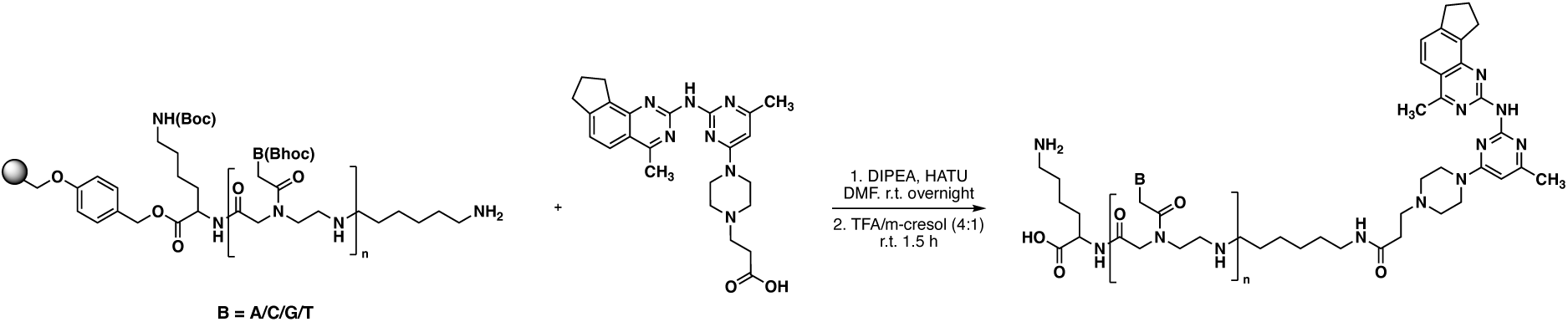
Synthesis of PNA GL-Os P6-P15 (n 6-15).

Finally, to probe the role of sequence complementarity, a series of triple-point mutation GL-Os were designed based on the 15 nt oligonucleotide. In these GL-Os, three consecutive bases were replaced with thymine, while any pre-existing thymine residues were substituted with cytidine to maintain overall base composition. The resulting variants (**GL-Os TM1-3**, **TM4-6**, **TM7-9**, **TM10-12**, and **TM13-15**) allow systematic evaluation of how mismatch position affects hybridization and target selectivity (see Table S2 for sequences and Table S3 for yields).

### Binding properties of the oligonucleotide modified GL-Os

The G4 target selected for this study was the *c-MYC* G4, known as Pu24T, a stabilized mutant of the native Pu27 sequence. This construct includes the G4-forming sequence along with a 20-nt 5’-flanking sequence complementary to the oligonucleotide in the GL-Os (Table S1). Throughout this study, the term G4 DNA refers specifically to the *c-MYC* Pu24T sequence along with this 5’-flanking sequence.

Binding of the GL-Os to the *c-MYC* G4 target was characterized by microscale thermophoresis (MST) and proton nuclear magnetic resonance (^1^H NMR) spectroscopy. MST provided dissociation constants (*K*_D_) from dose-response curves with fluorescently labelled G4 DNA (Table S1), while NMR monitored imino proton resonances to detect G4 interactions (10-12 ppm) and duplex formation (12-14 ppm). As controls, the G4-ligand alone and the individual unconjugated oligonucleotides were analyzed in both MST and NMR experiments (Figures 2, S1-S4, S6-S8,) Together, these complementary techniques allowed quantitative and structural assessment of GL-O binding.

**Figure 2.**
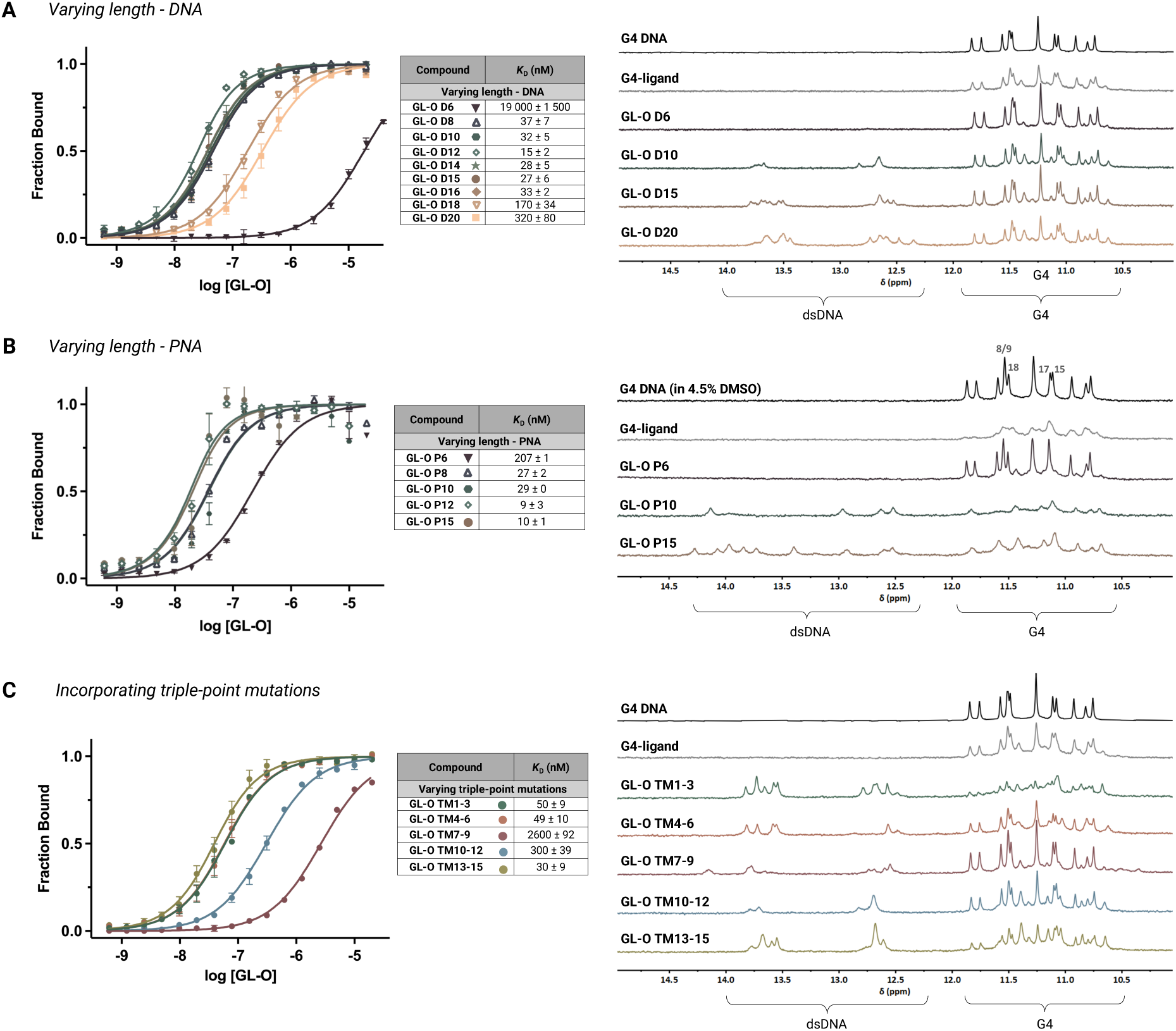
Binding affinity and interaction of the oligonucleotide modified GL-Os with the G4 DNA using MST to obtain dose-response curves and dissociation constants (*K*_D_) values (error bars correspond to two independent measurements). GL-Os were serial diluted to 5’-fluorescently labelled *c-MYC* Pu24T G4. ^1^H NMR profiles of the G4 DNA in 1:1 molar ratios with *c-MYC* Pu24T with complementary flanking sequence (90 μM). DNA-based constructs were analysed in aqueous buffer, whereas PNA-based constructs required the presence of 4.5% (v/v) DMSO for NMR measurements to ensure solubility; DMSO had no effect on MST-derived binding parameters. G4 imino proton signals appear between 10-12 ppm and double-stranded DNA imino proton signals between 12-14 ppm. A) Varying the length of the DNA oligonucleotide (**GL-Os D6**-**D20**). B) Varying the length of the PNA oligonucleotide (**P6-P15**). C) Incorporating triple-point mutations in the oligonucleotide sequence (**GL-Os TM1-3, TM4-6**, **TM7-9**, **TM10-12**, **TM13-15**).

#### Effect of oligonucleotide length

A clear relationship between oligonucleotide length and binding affinity was observed for the DNA GL-Os. MST analysis showed that conjugates with oligonucleotides shorter than 8-nt displayed negligible binding, indicating that 8-nt represents the minimal effective length required for hybridization under these conditions (Figure 2A). Within the range of 8-16 nt, the DNA based **GL-Os D8**-**D16** displayed comparable affinities, with *K*_D_ values ranging from 15 to 37 nM, suggesting similar binding efficiency within this range. Unexpectedly, longer oligonucleotides in the **GL-Os D18** and **D20** exhibited reduced binding affinity, with *K*_D_ values increasing to 170 and 320 nM, respectively. This reduction may reflect slower hybridization kinetics. Consistent with this interpretation, extended incubation (24 h) decreased the *K*_D_ values for **GL-Os D18** and **D20**, suggesting that full hybridization requires additional time for longer oligonucleotides in the GL-Os (Figure S5).

NMR spectroscopy corroborated these findings (Figure 2A). Increasing oligonucleotide length in GL-Os from 6-nt to 20-nt resulted in the appearance of imino proton signals between 12–14 ppm, corresponding to double-stranded DNA formation. The G4 imino proton signals between 10–12 ppm was retained for all GL-Os, although binding of the G4-ligand affected the G4 structure leading to the appearance of additional peaks in this region, consistent with previous reports.^10,11^ Extension beyond 10-nt did not further alter the G4-specific imino resonances, indicating that once a minimal hybridization interface is established, additional base pairs do not affect the interaction between the G4 and the G4-ligand. The shortest construct (**GL-O D6**) showed no detectable hybridization and minimal perturbation of the G4 imino signals, consistent with its weak binding observed by MST and the behavior of the unconjugated oligonucleotide (Figure S6).

#### Effect of backbone composition

MST analysis revealed that substitution of the DNA backbone with PNA enhanced binding affinity (Figure 2B). For equivalent lengths, the PNA-based GL-Os exhibited lower *K*_D_ values than their DNA counterparts. For instance, **GL-O P15** showed stronger binding affinity than **GL-O D15**. This increase in affinity is consistent with the neutral, peptide-like PNA backbone, which eliminates electrostatic repulsion between the oligonucleotide and the DNA target and thereby stabilizes the duplex.

NMR spectroscopy supported these findings (Figure 2B) and additionally revealed the lower aqueous solubility of the PNA constructs, which required an increased DMSO content in the NMR experiments (4.5%). This change in solubility gave minor alterations of the G4 imino protons with a separation of G-8/9 and G-18 and a merge of G-15 and G-17 (Figure 2B). The NMR showed that PNA-based GL-Os displayed more pronounced broadening and partial shifting of the G4 imino proton signals (10–12 ppm), compared with the DNA analogues, indicating stronger or more persistent interactions with the G4 structure. Signals corresponding to dsDNA regions (12–14 ppm) were also prominent, consistent with extensive hybridization. However, similar to the DNA **GL-O D6**, the shortest PNA **GL-O P6** failed to hybridize and showed only minimal effects on the G4 signals. Interestingly, the unconjugated PNA oligonucleotides alone induced small shifts in the G4 imino region (Figure S7), reflecting their inherent effect on the surrounding environment. However, when conjugated to the G4-ligand, the resulting GL-Os produced broader and more intense perturbations of the G4 signals, implying that the G4-ligand and the PNA act cooperatively to stabilize the complex. Furthermore, both **GL-Os P10** and **P15** gave strong broadening of the G4-signals, which was not evident from the binding affinity where **GL-O P15** had a significantly better affinity than **GL-O P10**.

#### Effect of sequence complementarity

For the triple-point mutation GL-O series (**GL-Os TM1-3** to **TM13-15**), MST results revealed a clear mismatch position dependence. Mutations positioned closer to the center of the oligonucleotide sequence (**GL-O TM7-9**) resulted in higher *K*_D_ values, indicating weaker binding, whereas mutations near the terminal regions (**GL-Os TM1-3** and **TM13-15**) had only minor effects (Figure 2C). NMR analysis supported these observations (Figure 2C). All mutation variants produced detectable dsDNA signals, but the degree of G4 interaction varied. Constructs with terminal mismatches exhibited extensive broadening and downfield shifts of the G4 imino resonances, indicating that the ligand remained engaged with the G4. In contrast, central mismatches (**GL-O TM7–9**) led to diminished broadening and weaker G4 signal intensity, consistent with reduced ligand binding when hybridization in the core region is disrupted. Interestingly, mismatches adjacent to the G4 (**GL-O TM1–3**) appeared to enhance the ligand’s effect on the G4 structure, possibly by introducing local flexibility while maintaining the spatial configuration necessary for optimal G4 interaction.

### G4-stabilization of the oligonucleotide modified GL-Os

The GL-O strategy has previously been shown to stabilize G4 DNA structures, as demonstrated by thermal stability assays and increased stalling with Taq DNA polymerase stop assays.^10,11^ Consistent with its binding mechanism, oligonucleotide hybridisation and G4-ligand interaction are expected to act synergistically to stabilize the G4 conformation. To assess this, thermal stability was first evaluated using ^1^H NMR by gradually increasing the temperature from 25 °C to 35 °C and then 45 °C, followed by 2.5 °C increments up to 65 °C (Figure 3A). The GL-Os in complex with the G4 DNA at a 1:1 molar ratios were compared to the G4 DNA and G4-ligand alone with and without DMSO (Figure S9). To determine whether thermal stabilization of the G4 structure results in a functional consequence, a Taq polymerase stop assay was performed (Figure 3B).

**Figure 3.**
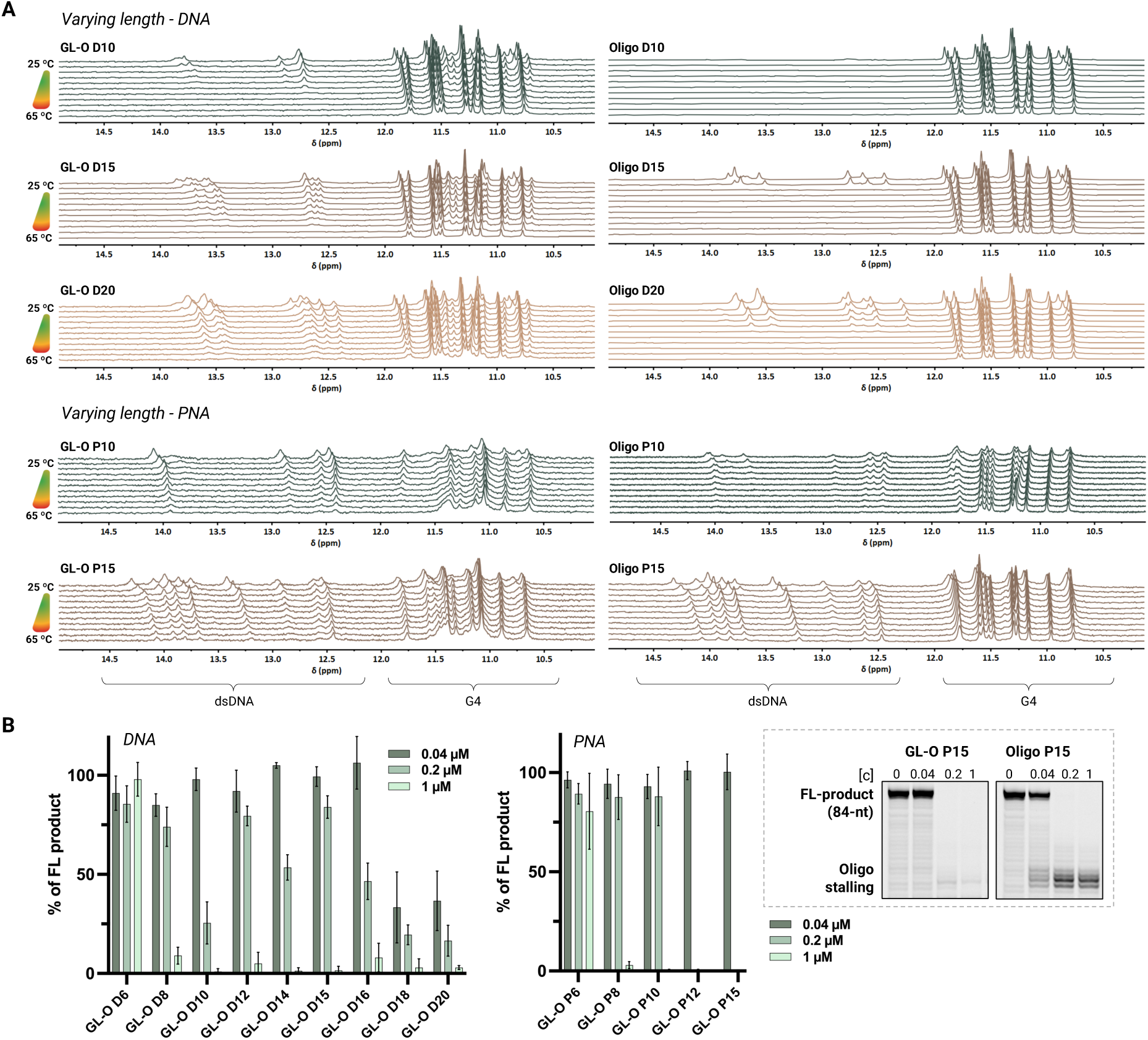
G4 stabilization of the *c-MYC* Pu24T G4 DNA structure by the modified GL-Os. A) Comparison between unconjugated oligonucleotides and with G4-ligand using ^1^H NMR **GL-Os** and **oligos D10**, **D15** and **D20** and **GL-Os** and **oligos P10** and **P15**. Spectra were recorded of *c-MYC* Pu24T with complementary 5’-flanking sequence at 25 and 35 °C and a 2.5 °C temperature ramp from 45-65 °C. Spectra for DNA-based constructs were recorded in aqueous buffer, whereas PNA-based constructs required the presence of 4.5% (v/v) DMSO to ensure solubility. G4 imino signals appear between 10-12 ppm and double-stranded DNA signals appear between 12-14 ppm. B) Taq Polymerase stop assay that measures stabilization of the G4 DNA in presence of the modified **GL-Os D6**-**D20** and **P6**-**P15**. Quantification of the Taq polymerase stop assay, shown as the full-length (FL, 84-nt) DNA product, is expressed as a % of the full-length band intensity that was observed in the control reaction containing only the G4 template. Data represent the mean ± standard deviation from three independent experiments.

The assay, previously described,^10,11^ evaluates DNA polymerase progression on a G4-contanining DNA template in the presence or absence of GL-Os. While Taq DNA polymerase exhibits baseline stalling at the G4 site, increase stalling, reflected by reduced full-length (FL) DNA product, indicates enhanced G4 stabilisation that impedes polymerase G4 bypass.

#### Effect of oligonucleotide length

Among the DNA-based GL-Os tested (**D10**, **D15**, and **D20)**, increasing oligonucleotide length correlated with elevated melting temperatures (*T*_m_) for both the duplex DNA and the G4-ligand dissociation from the G4 DNA structure (Figure 3A). The shortest construct, **GL-O D6**, failed to hybridize with the G4 flanking sequence and showed only minor G4-ligand induced effects, which was rapidly lost upon increasing the temperature (Figure S10). In contrast, the longest construct, **GL-O D20**, exhibited both the highest duplex *T*_m_ and the highest G4-ligand dissociation temperature (about 65 °C), suggesting that although hybridization kinetics are slower for longer oligonucleotides, the resulting complexes are more thermally stable. Interestingly, for all tested DNA GL-Os, the melting of the duplex occurred at approximately the same temperature as the loss of G4/G4-ligand interaction, indicating a coordinated stabilizing effect (Figure 3A). When compared with the corresponding unconjugated oligonucleotides, it is evident that conjugation of the G4-ligand in the GL-Os substantially increases duplex stability, clearly illustrated by the enhanced melting profile of **GL-O D20** relative to its unconjugated counterpart **oligo D20** (Figures 3A).

Consistent with these thermal stabilization trends (Figure 3A), all DNA GL-Os, except the shortest construct **GL-O D6,** showed a dose-dependent increase on polymerase stalling, indicative of enhanced G4-stabilization, (Figure 3B and S11). Although this assay does not resolve fine differences between oligonucleotide lengths, it reveals a general trend in which longer oligonucleotides, such as in **GL-Os D16** and **D18** induce stronger polymerase stalling, resulting in reduced formation of FL-product (Figure 3B). This correlates well with the high thermal stabilization induced by **GL-O D20** in the NMR melting experiments.

#### Effect of backbone composition

Overall, the PNA conjugates displayed markedly higher melting temperatures than the DNA-based GL-Os, indicating greater thermal stability of both the duplex and G4 interactions (Figure 3A). **GL-O P15** retained strong duplex and G4-ligand related signals even at the highest temperature measured (65 °C). However, unlike the DNA GL-Os, the unconjugated PNA oligonucleotides exhibited similar duplex melting profiles to their conjugated counterparts (Figure 3A). This reflects the inherently stronger hybridization of PNA, which produces highly potent GL-Os but also reduces their dependence on G4-ligand binding. This is a factor that should be considered in terms of potential nonspecific interactions. Furthermore, the melting behaviour of the PNA-based GL-Os differed from that of the DNA series and more closely resembled the profile of the free G4-ligand, likely reflecting the large difference in physicochemical properties between the PNA and DNA backbones.

Consistent with the NMR data, in the Taq polymerase stop assay, constructs containing longer PNA oligonucleotides exhibited more pronounced inhibition of polymerase progression, consistent with stronger G4 stabilization (Figure 3B and S12). The dose-response profiles for the PNA-based GL-Os displayed a steep transition, with a narrow range between concentrations showing no effect and those producing complete inhibition at very low concentrations. Both **GL-Os P12** and **P15** where inactive at 0.04 μM but induced a complete loss of full-length product at 0.2 μM (Figure 3B).

To further probe the strong affinity of PNA relative to DNA, we examined the corresponding unconjugated PNA oligonucleotides. In contrast to the DNA oligos, the PNA oligos strongly inhibited polymerase extension, but this effect did not originate from G4 stabilization. Instead, the data suggest that polymerase progression is blocked when encountering a PNA:DNA hybrid downstream of the G4. This was evident from **oligo P15** which showed a strong oligo stalling band (Figure 3B).

Importantly, this artifact was not observed for the PNA-based GL-Os. Upon conjugation of the G4 ligand, the GL-Os showed inhibition exclusively attributable to G4 stabilization, with no evidence of PNA-induced polymerase stalling (Figure 3B and S11). This indicates that the strong affinity of PNA is translated into G4 stabilization only when conjugated to the G4-ligand, an effect that was particularly pronounced for the longer 12- and 15-nt GL-Os. This conclusion was further supported by circular dichroism measurements, in which melting temperature profiles of G4 DNA alone were compared with those obtained in the presence of the G4-ligand, unconjugated PNA oligonucleotides (**P6, P10, P15**), or the corresponding GL-Os (**P6, P10, P15**) (Figure S12). While the PNA oligonucleotides alone did not stabilize the G4 DNA, ligand-conjugated PNA GL-Os produced substantial G4 stabilization, exceeding that of the G4-ligand alone, particularly for **GL-O P10** and **GL-O P15**. This further support that the GL-O design preserves G4-specific activity while mitigating the nonspecific hybridization-driven interference observed for unconjugated PNA oligonucleotides.

#### Effect of sequence complementarity

GL-Os containing central triple-point mutations (GL-Os **TM4-6**, **TM7-9** and **TM-10-12)**, which exhibited reduced binding affinity, also failed to induce thermal stabilization effects (Figure S13). These constructs showed minimal or no interaction with the G4 even at 35 °C. In contrast, GL-Os with terminal mutations retained G4 association up to 50-55 °C, consistent with the higher tolerance for mismatches near the duplex ends observed in the binding assays (Figure 3A). For the unconjugated oligonucleotides, no G4 stabilization was observed (Figure S13).

Consistent with these observations, in the Taq polymerase stop assay, no significant difference in FL product were observed at 0.04 μM as all triple-mutant GLOs displayed minimal effects at this concentration. However, at increasing concentrations, a clear trend emerged where GL-Os with central mutations (**TM7-9** and **TM10-12**) exhibited the weakest G4 stabilization, yielding approximately 50% FL product at the highest concertation tested (Figure S14), consistent with the NMR melting data.

### Stability of GL-Os toward endonuclease

GL-Os exhibit strong affinity and robust G4 stabilization *in vitro*, but their suitability for cellular applications also depends on nuclease resistance. To assess this, we incubated DNA- and PNA-based GL-Os with the length of 15-nt, together with their unconjugated oligonucleotide as controls, in the presence of internally cleaving nuclease (Figure 4).

**Figure 4.**
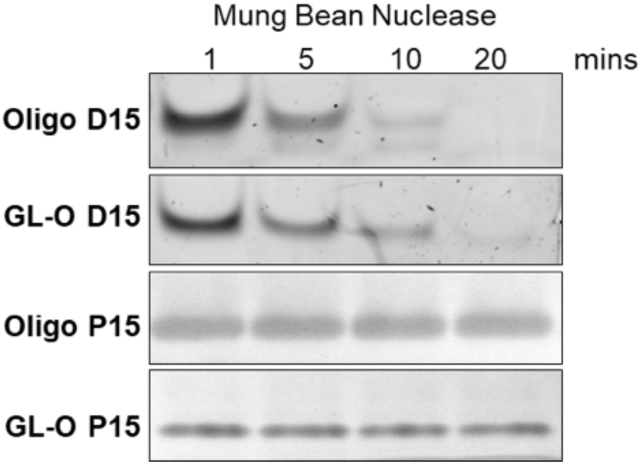
*In vitro* digestion of oligonucleotides by Mung Bean nuclease. Unconjugated and Conjugated Oligos were incubated with Mung bean nuclease at 30 °C and quenched at the indicated time points. Digestion products were analyzed by PAGE - DNA on 16% TBE gels stained with GelRed and PNA on 4–20% SDS–PAGE gels stained with Instant Blue. Gels were imaged using a ChemiDoc (Bio-Rad) system.

Time-dependent degradation of DNA oligonucleotides was observed upon incubation with a restriction endonuclease Mung Bean nuclease, as evidenced by progressive loss of full-length DNA oligonucleotides (unconjugated and conjugated) on TBE PAGE. Interestingly, the conjugated **GL-O D15** showed to be more metabolically stable than its unconjugated **oligo D15** (Figure 4).

In contrast, PNA oligonucleotides (unconjugated and conjugated) remained intact over the same incubation period, and no detectable degradation products are observed on SDS PAGE, indicating resistance to MBN digestion (Figure 4), highlighting that PNA improves the metabolic stability of the GL-O, a feature required for cellular activity.

## Discussion

This study provides a systematic analysis of the molecular determinants governing G4 recognition and stabilization by G4-ligand–conjugated oligonucleotides (GL-Os). By varying oligonucleotide length, backbone composition, and sequence complementarity, we identify the key parameters that control GL-O binding affinity, structural engagement, and functional G4 stabilization.

A central component of this work was the successful synthesis and characterization of a comprehensive GL-O library, comprising nineteen conjugates alongside their unconjugated counterparts. The use of complementary conjugation strategies, including a robust copper-free click reaction for DNA-based GL-Os and solid-phase amide coupling for PNA-based constructs, enabled the modular construction of structurally diverse GL-Os. This dual synthetic approach allowed incorporation of chemically distinct oligonucleotide backbones without altering the G4-ligand scaffold, allowing systematic investigation of how oligonucleotide length, backbone composition, and sequence complementarity influence G4 targeting and stabilization.

Genome-wide alignment analyses across three representative G4 targets (c-MYC, K-Ras, and Helicase B) established an important conceptual framework for GL-O selectivity. These analyses revealed that sequence uniqueness is highly target-dependent, with the c-MYC Pu27 region retaining single-locus specificity at oligonucleotide lengths as short as 14 nucleotides, whereas other loci require longer sequences to achieve comparable discrimination. Although shorter oligonucleotides showed partial complementarity to additional genomic loci, such matches are unlikely to result in functional off-target effects. The GL-O strategy relies on a dual-binding requirement, whereby productive engagement necessitates both oligonucleotide hybridization to a flanking single-stranded region and simultaneous binding of the G4-ligand to a neighboring G4 structure. This interdependence implies that the oligonucleotide does not need to achieve full genome-wide uniqueness, as functional binding will occur only where sequence complementarity coincides with a compatible G4 motif. Thus, the dual-recognition mechanism provides an inherent safeguard against nonspecific interactions while allowing flexibility in oligonucleotide design.

Experimental characterization further demonstrate that oligonucleotide length exerts a dual effect on GL-O performance. Short guide strands (< 8 nt) fail to hybridize effectively and show minimal G4 interaction, while intermediate lengths (10–16 nt) bind efficiently but yield moderate stabilization. In contrast, longer oligonucleotides (≥ 18 nt) display slower hybridization kinetics yet produce markedly higher thermal and polymerase-mediated G4 stabilization. This indicates that although longer guides associate more slowly, they form more persistent and thermally stable complexes once bound. The stabilization data for the DNA GL-Os further reveal a strong interdependence between duplex formation and G4-ligand interaction, demonstrating that both elements act cooperatively to reinforce each other. This dual engagement not only enhances overall complex stability but also anchors the guide oligonucleotide at the correct genomic site, thereby minimizing the likelihood of off-target binding.

Backbone composition further modulates these effects. PNA-based GL-Os exhibited greater thermal stability and prolonged G4 association relative to their DNA counterparts, consistent with the high affinity of the neutral PNA backbone. Importantly, G4-ligand conjugation transforms PNA guides from nonspecific polymerase-blocking agents into selective G4 stabilizers, thereby preserving G4-specific activity and mitigating hybridization-driven off-target effects. Moreover, PNA GL-Os exhibit a propensity to enhance the metabolic stability of GL-Os, a critical prerequisite for sustaining subsequent cellular activity. This property underscores the advantageous nature of PNA GL-Os within the GL-O strategy.

Sequence complementarity emerged as another critical determinant of GL-O function. Central mismatches within the oligonucleotide severely compromised binding affinity and G4 stabilization, whereas terminal mismatches were better tolerated. This behavior is consistent with general thermodynamic principles of nucleic acid pairing, where internal mismatches impose the greatest destabilizing penalties, and demonstrates that central complementarity is essential not only for duplex formation but also for maintaining effective G4-ligand engagement.

Together, these findings show that effective GL-O binding and G4 stabilization depend on a balance between oligonucleotide hybridization and G4-ligand engagement. Oligonucleotide length, backbone composition, and sequence complementarity jointly determine this balance: short sequences hybridize inefficiently, longer sequences hybridize more slowly but form more stable complexes, and central mismatches destabilize duplex formation, while PNA substitution enhances binding strength but requires ligand conjugation to maintain specificity. This interdependence stabilizes the oligonucleotide at the target site and limits off-target interactions. Overall, the results underscore the pivotal role of the oligonucleotide component in defining GL-O behaviour and highlight the modular, chemically adaptable nature of the strategy, enabling rational design of GL-Os for probing G4 biology and developing targeted therapeutics, particularly in oncogenic contexts such as the c-MYC promoter.

## Material and methods

### Conjugation of G4-ligand with DNA oligonucleotides

5′-Hexylamine oligonucleotide (50 nmol) was dissolved in 0.1 M NaHCO₃ (50 µL). BCN-NHS ester (20 equiv., 0.2 M in DMSO) was added, and the mixture was shaken for 24 h. The product was precipitated twice using 3 M NaOAc and 4x volume ethanol, washed with ethanol, dried under N₂, and redissolved in 0.1 M NaHCO₃. An azide-functionalized G4-ligand (2.5 equiv., 10 mM in DMSO) was added, and the reaction was shaken for 24 h. The mixture was diluted, filtered, and purified by RP-HPLC (C18 column, 2 mL/min, 5–60% ACN in 50 mM TEAA buffer). Conjugates were confirmed by HRMS (ESI⁻).

### Peptide nucleic acid synthesis

PNA synthesis was done on an automated solid phase peptide synthesizer (SPPS) on a 20 µmol scale using Fmoc-Lys(Boc)-Wang (0.61 mmol/g loading) resin for **P6**-**P10** and Fmoc-Lys(Boc)-Wang (0.2 - 0.4 mmol/g) for **P12**-**P15**. The resin was allowed to swell in DCM for 30 min. The Fmoc protecting group was removed by treating the resin with 20% (v/v) piperidine in DMF. Fmoc-PNA(Bhoc)-OH monomers (0.50 M in DMF) were coupled using HATU (0.49 M in NMP) and DIPEA (2.0 M in DMF) for 60 minutes. A capping step was performed after each coupling using 50% (v/v) Ac_2_O in DCM for 40 minutes. Cleavage and deprotection were performed using a mixture of TFA:m-cresol (4:1) for 1 h at room temperature. The cleavage mixture was filtered into ice-cold diethyl ether (Et^2^O), and the resulting precipitate was diluted, filtered, and subjected to purification by RP-HPLC (C18 column, 2 mL/min, 0-30% then 30-100% solvent B (100% ACN) and solvent A (0.1% FA). The conjugated products were confirmed by HRMS (ESI^+^).

### Conjugation of G4-ligand with PNA oligonucleotides

The G4-ligand (0.25 M, 2.5 eq) was solubilized in NMP and added to the resin (5 µmol) along with HATU (0.24 M in NMP) and DIPEA (1.0 M in DMF). The reaction was allowed to shake for 24 h at room temperature. The conjugated PNAs were cleaved from the resin following the aforementioned procedures.

### Microscale thermophoresis (MST)

Cy5-labeled G4 DNA was annealed in MST buffer (10 mM potassium phosphate, 100 mM KCl, 0.05% Tween 20, pH 7.4) by heating at 96 °C for 5 min and cooling to room temperature. MST measurements were performed on a Monolith NT.115 using standard capillaries, with G4 DNA at 20 nM and serially diluted GL-Os, unconjugated oligonucleotides and G4-ligand (5-40 µM). DNA-based oligonucleotides were prepared in aqueous buffer. For PNA-based GL-Os and unconjugated PNA oligonucleotide controls, experiments were performed both in the presence and absence of DMSO, and no differences in thermophoresis traces or derived binding affinities were observed. Dissociation constants (*K*^D^) were obtained using Monolith analysis software and dose-repones curves were plotted in GraphPad Prism 10.

### Proton nuclear magnetic resonance spectroscopy (^1^H NMR)

G4 DNA (110 μM) was annealed in 10 mM potassium phosphate buffer (3 mM KCl, pH 7.4) following same procedure as for MST. Before measuring, 10% D₂O was added, and the sample (100 μM) was transferred to a 3 mm tube. GL-O, unconjugated oligonucleotide (1 mM), or G4-ligand (10 mM) were added in separate experiments to achieve a 1:1 molar ratio relative to G4 DNA, and ¹H NMR spectra were recorded after equilibration. DNA-based GL-Os were prepared in aqueous buffer, whereas PNA-based constructs required the addition of DMSO to a final concentration of 4.5% (v/v) to ensure complete solubility. Experiments were performed on an 850 MHz Bruker AVANCE III HD spectrometer at 298 K using excitation sculpting (512 scans). Thermal melting profiles were recorded at 308 K then a temperature ramp of 2.5 K from 318 to 338 K. Data were processed with MestreNova 10.0.2.

### Taq polymerase STOP assay

DNA templates were annealed with a fluorescent primer in 100 mM KCl by heating to 95 °C and cooling to room temperature. Reactions (40 nM template) were prepared in 1×Taq Buffer (10 mM Tris-HCl pH 8.8, 50 mM KCl), 1.5 mM MgCl_2_, 0.05 U/μL Taq Polymerase, and compound as indicated. DNA-based oligonucleotides were prepared in water. For PNA-based GL-Os and unconjugated PNA oligonucleotide controls, assays were performed both in the presence and absence of DMSO, and no differences in polymerase stalling or full-length product formation were observed. After 10 min on ice, dNTPs (100 μM) were added and samples incubated at 37 °C for 15 min. Reactions were stopped with 2×stop solution and separated on a 12% denaturing TBE gel. Fluorescence was detected using a Typhoon Scanner, and band intensities were quantified with ImageQuant TL 10.2.

### *In vitro* endonuclease digestion

Stability of oligos against Mung bean nuclease digestion was assessed. 4 μM oligos were incubated with 3 units of enzyme (MBN, NEB) in 1X MBN buffer (MBN) at 30 degrees. At indicated time points, reactions were quenched by adding DNA Gel loading dye (50 mM EDTA, 50 mM HEPES, 30% Glycerol, 0.001% Bromophenol Blue) and heating at 95 °C. Digestion products were analysed by polyacrylamide gel electrophoresis (PAGE). DNA samples were resolved on 16% TBE gels for 1 h at 100 V in 1X TBE buffer. PNA samples were loaded on 4-20% SDS PAGE gels (Mini-PROTEAN TGX Gels, Bio-Rad) and run for 1 h at 120 V in 1X SDS running buffer (Tris Glycine). TBE gels were stained with Gel red (Biotium), while SDS PAGE gels were stained with Instant Blue (Abcam). Gels were imaged using a ChemiDoc Imaging system (Bio-Rad).

### CD melting temperature assay

G4 (3 µM) was folded in K-phosphate buffer (10 mM, pH 7.4) containing KCl (3 mM) by heating for 5 minutes at 95 °C, then allowed to cool to room temperature. PNA in DMSO was added in a 1:1 ratio, resulting in a final DMSO concentration of 0.3%. CD measurements were performed on a J-1700 Circular Dichroism Spectrophotometer (Jasco International Co. Ltd.). A quartz cuvette with a path length of 1 mm was used for measurements. CD spectra were recorded between λ = 200-350 nm with an interval of 0.5 nm and a scan rate of 100 nm/min. Thermal melting was recorded between 25°C and 95°C with increments of 10°C at a speed of 1 °C/min.

### Sequence alignment analysis

The specificity of 5′ G4 flanking sequences of varying lengths was assessed by aligning these DNA sequences to the human reference genome. The human reference genome (GRCh38) was retrieved from the UCSC Genome Browser,^17^ and flanking sequence was obtained from NCBI nucleotide database. FASTA datasets containing flanking sequences of 5–20 bases were generated and each dataset was independently aligned to the human reference genome using Bowtie 2.0,^18^ with no mismatch allowed. The aligned reads in SAM files were converted to BAM format and subsequently sorted and indexed using SAMtools^19^ with default settings. The final BAM files were used for downstream sequence annotation and comparative analysis of mapping specificity across different flanking lengths.

## Data availability

Detailed synthetic procedures and conjugation methods, experimental data for the assays are provided in the ESI.

## Acknowledgements

Work in the Chorell lab was supported by grants from the Swedish Research Council (VR-NT 2021-04805), the Swedish Cancer Foundation (23 2793 Pj), Lion’s Cancer Research Foundation in Northern Sweden (LP 24-2352), and the Kempe Foundations (JCK-3159). We thank the Knut and Alice Wallenberg foundation program “NMR for Life” for NMR spectroscopy support. Work in the Wanrooij lab was supported by the Knut and Alice Wallenberg Foundation, Kempe Foundations (SMK21-0059), and the Swedish Research Council (VR-MH 2023-02160).

## Author contribution

E.C. and S.W. initiated and directed the project. A.A., S.K., S.W., E.C., S.V., A.B., K.A., N.C., C.K. and L.S. developed the project. A.A., E.C., and S.W. wrote the manuscript. C.K. performed the sequence analysis. Oligonucleotide synthesis was performed by T.B. and A.D. PNA and PNA GL-O synthesis was performed by S.K. and S.V.. Bioconjugations and biophysical assays were performed by A.A., S.K., and L. S.. Polymerase stop assays were done by A.B., K.A., and L.S. *In vitro* digestion of GL-Os was performed by N.C. and A.A.. Analysis of the data was done by all authors. All authors read and edited the manuscript.

## Declaration of interests

The authors declare no conflict of interest.

## Notes

### Competing Interest Statement

The authors have declared no competing interest.

## References

1. Dell’Oca, M.C., Quadri, R., Bernini, G.M., Menin, L., Grasso, L., Rondelli, D., Yazici, O., Sertic, S., Marini, F., Pellicioli, A., et al. (2024). Spotlight on G-Quadruplexes: From Structure and Modulation to Physiological and Pathological Roles. Int. J. Mol. Sci. 25, 3162.

2. Lane, A.N., Chaires, J.B., Gray, R.D., and Trent, J.O. (2008). Stability and kinetics of G-quadruplex structures. Nucleic Acids Res. 36, 5482–5515.

3. Hänsel-Hertsch, R., Beraldi, D., Lensing, S.V., Marsico, G., Zyner, K., Parry, A., Di Antonio, M., Pike, J., Kimura, H., Narita, M., et al. (2016). G-quadruplex structures mark human regulatory chromatin. Nat. Genet. 48, 1267–1272.

4. Lago, S., Nadai, M., Cernilogar, F.M., Kazerani, M., Domíniguez Moreno, H., Schotta, G., and Richter, S.N. (2021). Promoter G-quadruplexes and transcription factors cooperate to shape the cell type-specific transcriptome. Nat. Commun. 12, 3885.

5. Siddiqui-Jain, A., Grand, C.L., Bearss, D.J., and Hurley, L.H. (2002). Direct evidence for a G-quadruplex in a promoter region and its targeting with a small molecule to repress c-*MYC* transcription. Proc. Natl. Acad. Sci. 99, 11593–11598.

6. Chaudhuri, R., Bhattacharya, S., Dash, J., and Bhattacharya, S. (2021). Recent Update on Targeting c-MYC G-Quadruplexes by Small Molecules for Anticancer Therapeutics. J. Med. Chem. 64, 42–70.

7. Boddupally, P.V.L., Hahn, S., Beman, C., De, B., Brooks, T.A., Gokhale, V., and Hurley, L.H. (2012). Anticancer Activity and Cellular Repression of c-MYC by the G-Quadruplex-Stabilizing 11-Piperazinylquindoline Is Not Dependent on Direct Targeting of the G-Quadruplex in the c-MYC Promoter. J. Med. Chem. 55, 6076–6086.

8. Nuccio, S.P., Cadoni, E., Nikoloudaki, R., Galli, S., Ler, A.-J., Sanchez-Cabanillas, C., Maher, T.E., Fan, E., Guneri, D., Flint, G., et al. (2025). Chemically modified CRISPR-Cas9 enables targeting of individual G-quadruplex and i-motif structures, revealing ligand-dependent transcriptional perturbation. Nat. Commun. 17, 385.

9. Chambers, V.S., Marsico, G., Boutell, J.M., Di Antonio, M., Smith, G.P., and Balasubramanian, S. (2015). High-throughput sequencing of DNA G-quadruplex structures in the human genome. Nat. Biotechnol. 33, 877–881.

10. Berner, A., Das, R.N., Bhuma, N., Golebiewska, J., Abrahamsson, A., Andréasson, M., Chaudhari, N., Doimo, M., Bose, P.P., Chand, K., et al. (2024). G4-Ligand-Conjugated Oligonucleotides Mediate Selective Binding and Stabilization of Individual G4 DNA Structures. J. Am. Chem. Soc. 146, 6926–6935.

11. Abrahamsson, A., Berner, A., Golebiewska-Pikula, J., Chaudhari, N., Keskitalo, E., Lindgren, C., Chmielewski, M.K., Wanrooij, S., and Chorell, E. (2025). Linker Design Principles for the Precision Targeting of Oncogenic G-Quadruplex DNA with G4-Ligand-Conjugated Oligonucleotides. Bioconjugate Chem. 36, 724–736

12. Abrahamsson, A., Khwaja, S., Berner, A., Nath Das, R., Aasumets, K., Chaudhari, R., Wanrooij, S., and Chorell, E. (2026). Tuning Potency for Precision: The Role of the G4-Ligand in G4-Ligand–Conjugated Oligonucleotides Targeting Individual G-Quadruplex DNA Structures [Revised for RSC Chemical Biology] *Department of Chemistry and Department of Medical Biochemistry and Biophysics, Umea University.*

13. Sun, D., and Hurley, L.H. (2009). The Importance of Negative Superhelicity in Inducing the Formation of G-Quadruplex and i-Motif Structures in the c-Myc Promoter: Implications for Drug Targeting and Control of Gene Expression. J. Med. Chem. 52, 2863–2874.

14. Egholm, M., Buchardt, O., Nielsen, P.E., and Berg, R.H. (1992). Peptide nucleic acids (PNA). Oligonucleotide analogs with an achiral peptide backbone. J. Am. Chem. Soc. 114, 1895–1897.

15. Wan, W.B., and Seth, P.P. (2016). The Medicinal Chemistry of Therapeutic Oligonucleotides. J. Med. Chem. 59, 9645–9667.

16. Nielsen, P.E. (2002). Peptide Nucleic Acids - Methods and Protocols (Humana Press Inc.).

17. Casper, J., Speir, M.L., Raney, B.J., Perez, G., Nassar, L.R., Lee, C.M., Hinrichs, A.S., Gonzalez, J.N., Fischer, C., Diekhans, M., et al. (2025). The UCSC Genome Browser database: 2026 update. Nucleic Acids Research 54, D1331–D1335.

18. Langmead, B., and Salzberg, S.L. (2012). Fast gapped-read alignment with Bowtie 2. Nature Methods 9, 357–359.

19. Li, H., Handsaker, B., Wysoker, A., Fennell, T., Ruan, J., Homer, N., Marth, G., Abecasis, G., Durbin, R., and Subgroup, G.P.D.P. (2009). The Sequence Alignment/Map format and SAMtools. Bioinformatics 25, 2078–2079.

