## Supporting Information for "Dual Recognition Drives Site-Directed G-Quadruplex Stabilization: Exploring Oligonucleotide Design in G4 Ligand-Oligonucleotide Conjugates"

**Table S1. G4 DNA templates**

| Used in | G4 DNA | Sequence (5´-3´) |
|---|---|---|
| MST assay | *c-MYC* Pu24T | [Cy5]-AGT CAC TGA ATT GGA CGT GA TGA GGG TGG TGA GGG TGG GGA AGG |
| NMR Assay | *c-MYC* Pu24T | AGT CAC TGA ATT GGA CGT GA TGA GGG TGG TGA GGG TGG GGA AGG |
| Taq Polymerase Stop Assay | c-MYC Pu24T | AGT CA CTG AAT TGG ACG TGA TGA GGG TGG TGA GGG TGG GGA AGG CAC GTG AGT TGA GTG GAG TTG GAA GTA GGC ATA CCC CTA T |
| Polymerase Stop assay | 15-nt TET primer | [TET]-GCC TAC TTC CAA CTC |
| Polymerase Stop assay | 25-nt TET primer | [TET]-ATA GGG GTA TGC CTA CTT CCA ACT C |

G4 forming sequence (red), nucleotide gap (green), primer site (blue)

**Table S2. GL-O conjugates synthesized for this study**

| GL-O name | Conjugation method | Oligo 5´modification | Sequence length | Sequence (5´-3´) |
|---|---|---|---|---|
| **GL-O D6** | SPAAC | Amine-C6 | 6-nt | *TmCAmCGT |
| **GL-O D8** | SPAAC | Amine-C6 | 8-nt | *TmCAmCGTmCmC |
| **GL-O D10** | SPAAC | Amine-C6 | 10-nt | *TmCAmCGTmCmCAA |
| **GL-O D12** | SPAAC | Amine-C6 | 12-nt | *TmCACGTmCmCAATT |
| **GL-O D14** | SPAAC | Amine-C6 | 14-nt | *TmCAmCGTmCmCAATTmCA |
| **GL-O D15** | SPAAC | Amine-C6 | 15-nt | *TmCAmCGTmCmCAATTmCAG |
| **GL-O D16** | SPAAC | Amine-C6 | 16-nt | *TmCAmCGTmCmCAATTmCAGT |
| **GL-O D18** | SPAAC | Amine-C6 | 18-nt | *TmCAmCGTmCmCAATTmCAGTGA |
| **GL-O D20** | SPAAC | Amine-C6 | 20-nt | *TmCAmCGTmCmCAATTmCAGTGAmCT |
| **GL-O P6** | Amide coupling | Amine-C6 | 6-nt | TCACGT |
| **GL-O P8** | Amide coupling | Amine-C6 | 8-nt | TCACGTCC |
| **GL-O P10** | Amide coupling | Amine-C6 | 10-nt | TCACGTCCAA |
| **GL-O P12** | Amide coupling | Amine-C6 | 12-nt | TCACGTCCAATT |
| **GL-O P15** | Amide coupling | Amine-C6 | 15-nt | TCACGTCCAATTCAG |
| **GL-O TM1-3** | SPAAC | Amine-C6 | 15-nt | TmCAmCGTmCmCAATTmCAG |
| **GL-O TM4-6** | SPAAC | Amine-C6 | 15-nt | TmCAmCGTmCmCAATTmCAG |
| **GL-O TM7-9** | SPAAC | Amine-C6 | 15-nt | TmCAmCGTmCmCAATTmCAG |
| **GL-O TM10-12** | SPAAC | Amine-C6 | 15-nt | TmCAmCGTmCmCAATTmCAG |
| **GL-O TM13-15** | SPAAC | Amine-C6 | 15-nt | TmCAmCGTmCmCAATTmCAG |

mC stands for methylated cytidine. * Is phosphorothioate modification

### *Chromatograms and data of the GL-O conjugates*

### GL-O D6

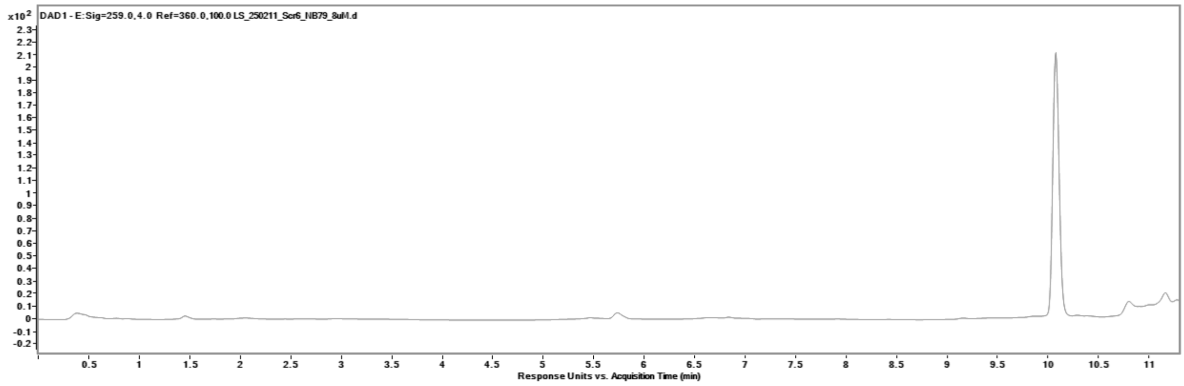

### GL-O D8

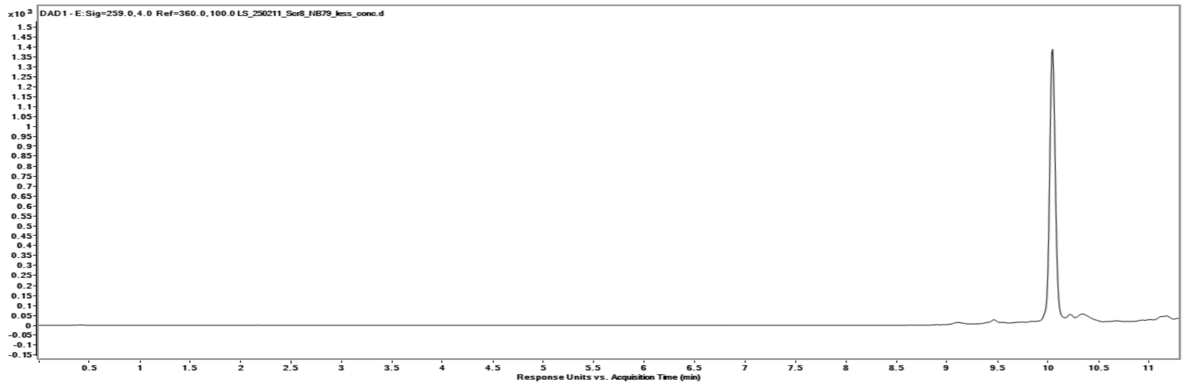

### GL-O D10

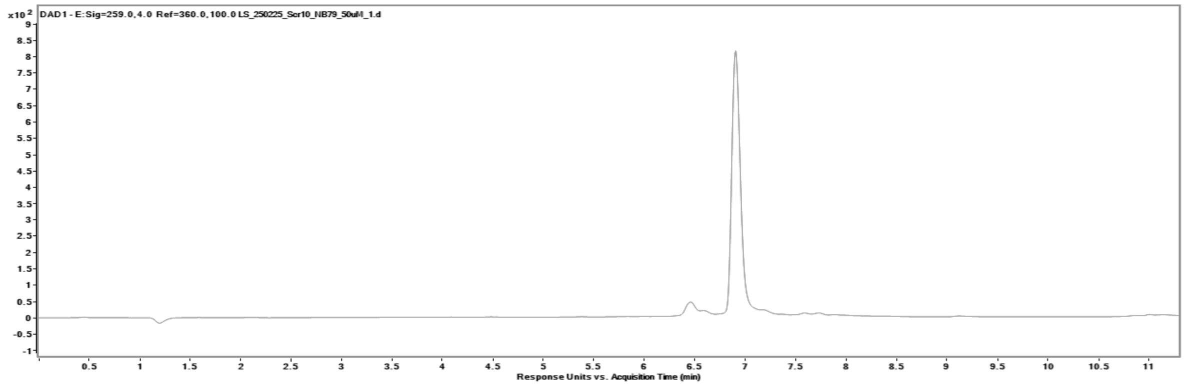

**GL-O D12**

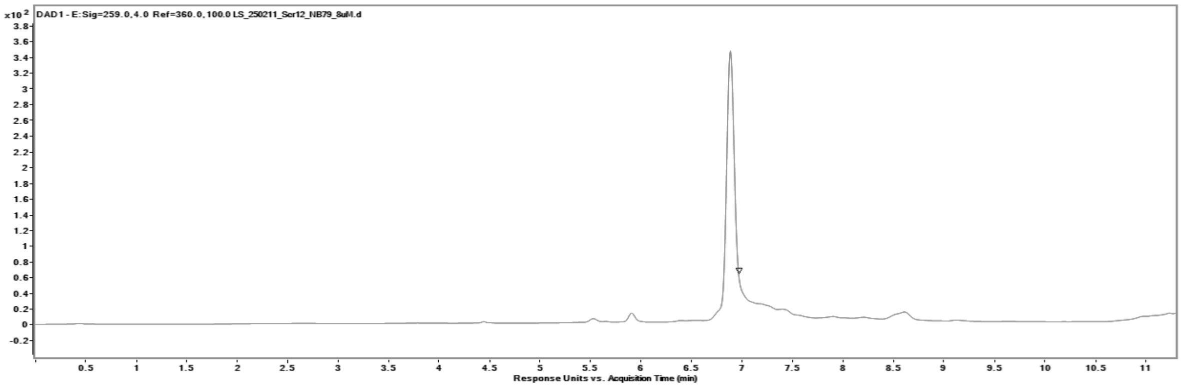

**GL-O D14**

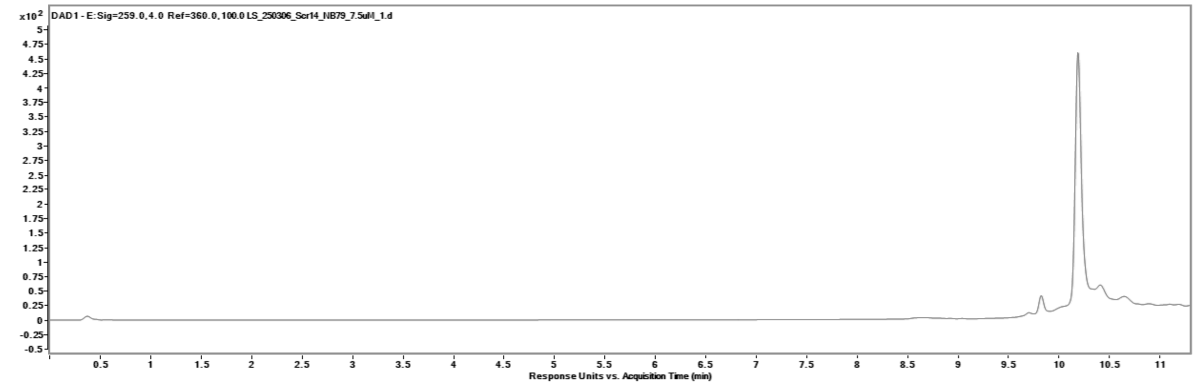

**GL-O D15**

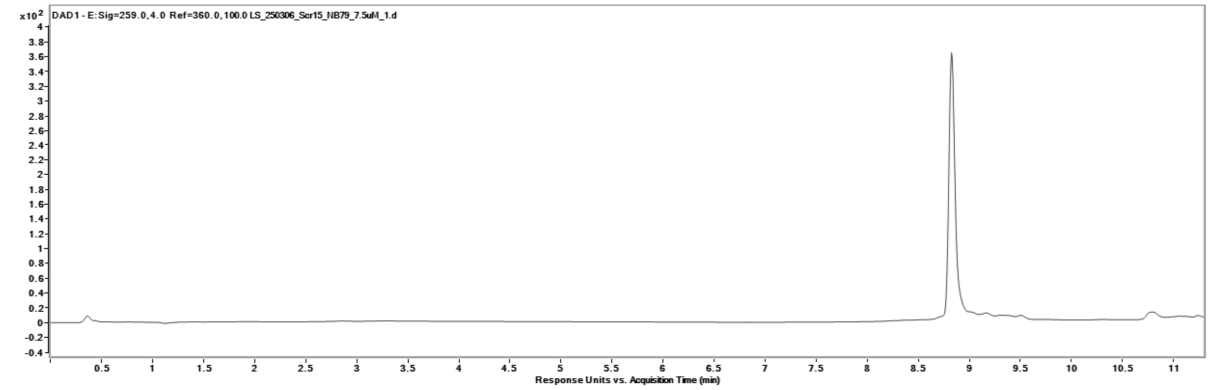

**GL-O D16**

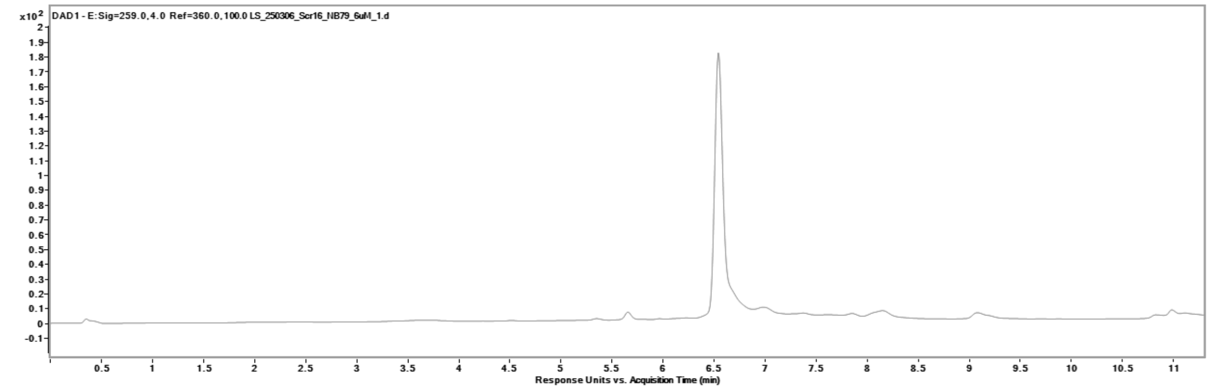

**GL-O D18**

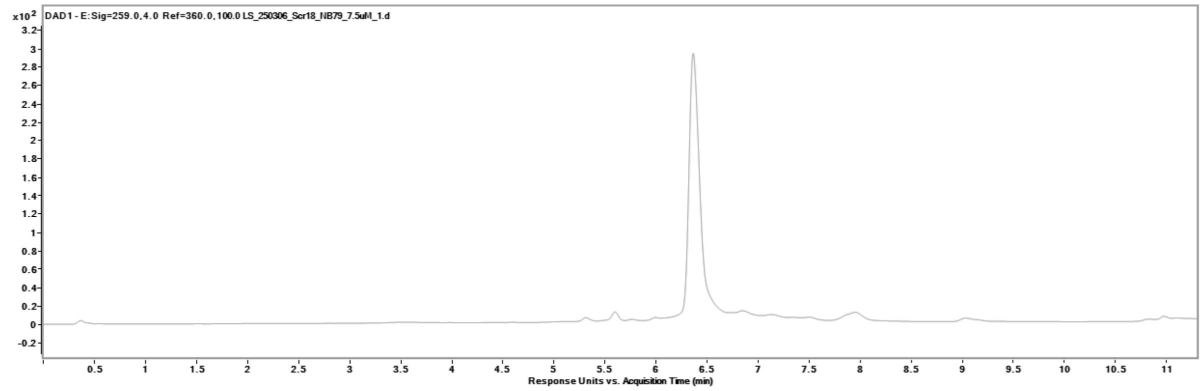

**GL-O D20**

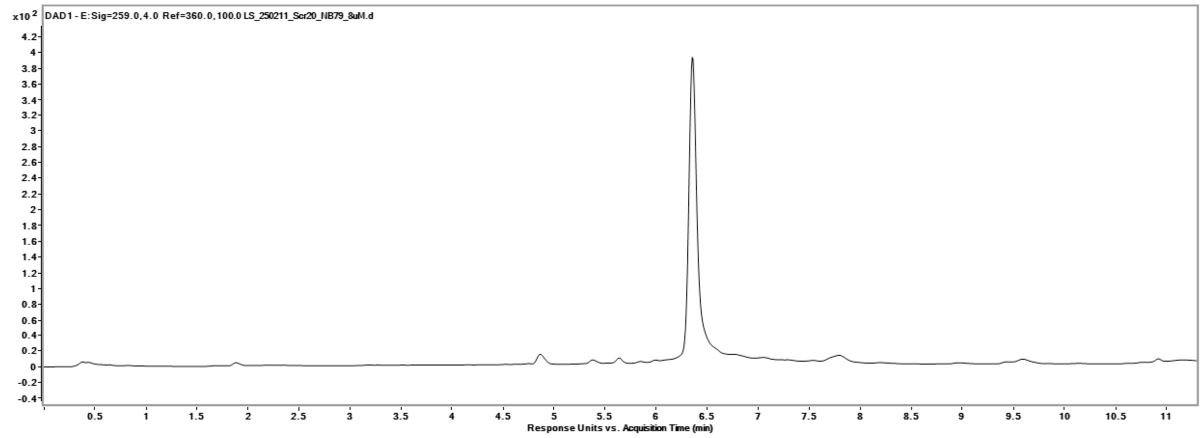

**GL-O P6**

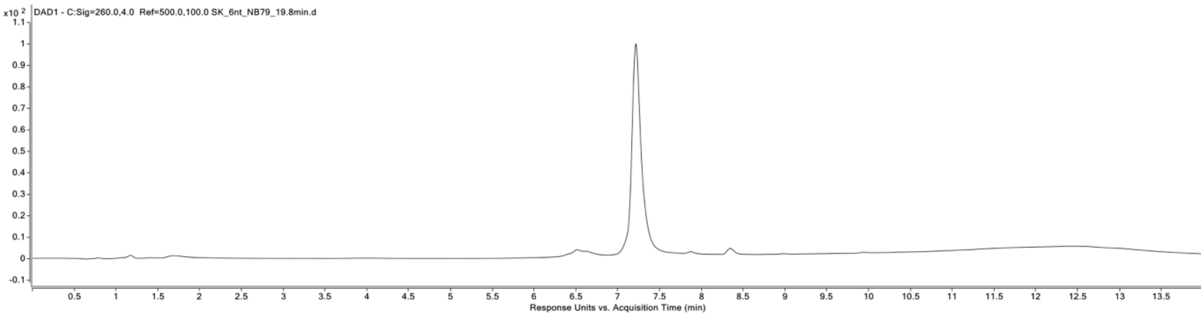

**GL-O P8**

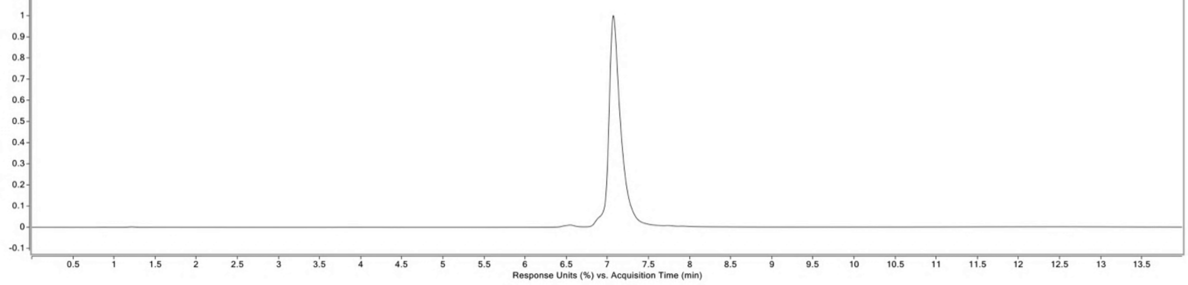

**GL-O P10**

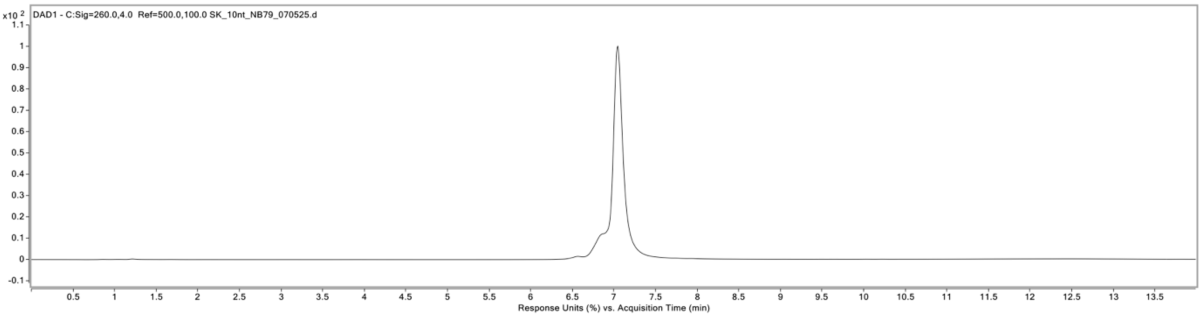

**GL-O P12**

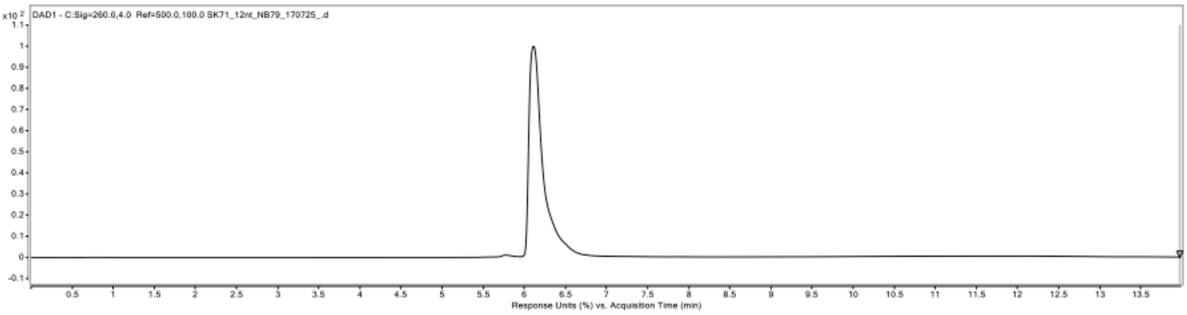

**GL-O P15**

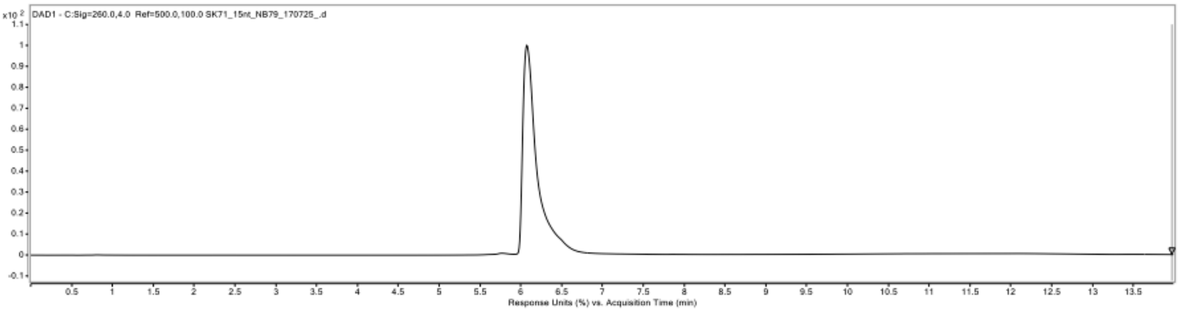

**GL-O TM1-3**

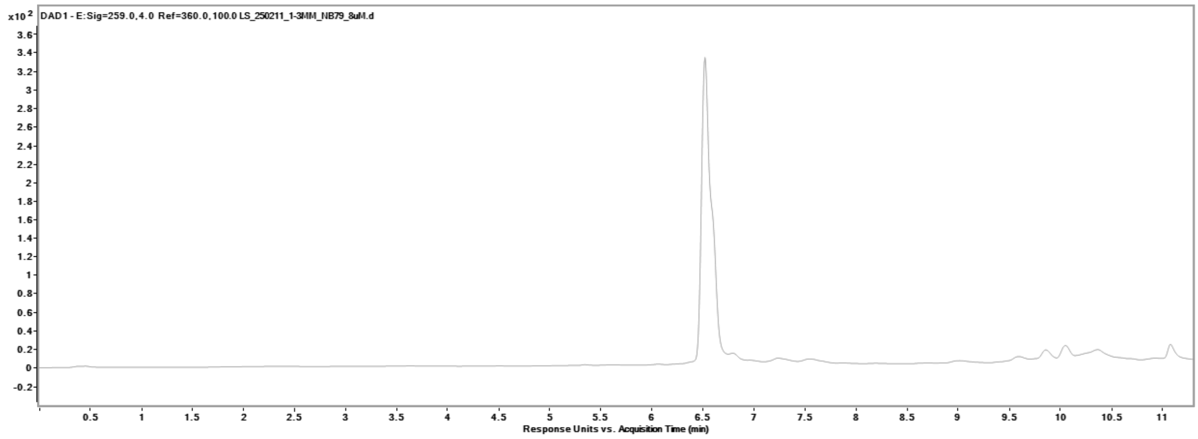

**GL-O TM4-6**

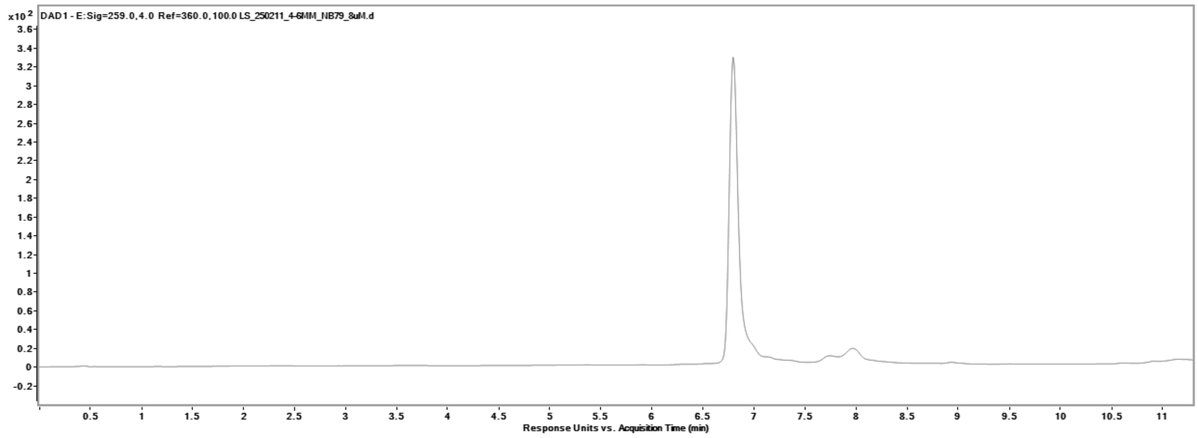

**GL-O TM7-9**

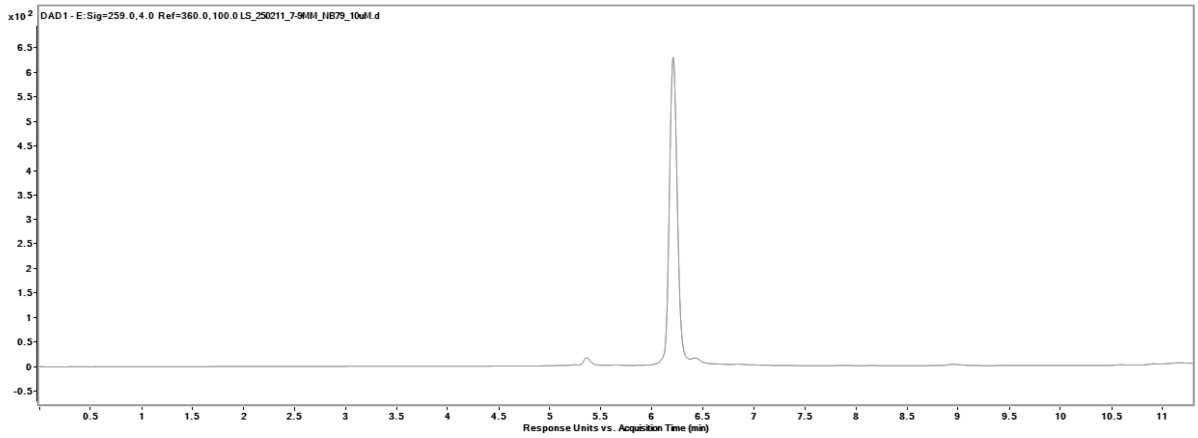

**GL-O TM10-12**

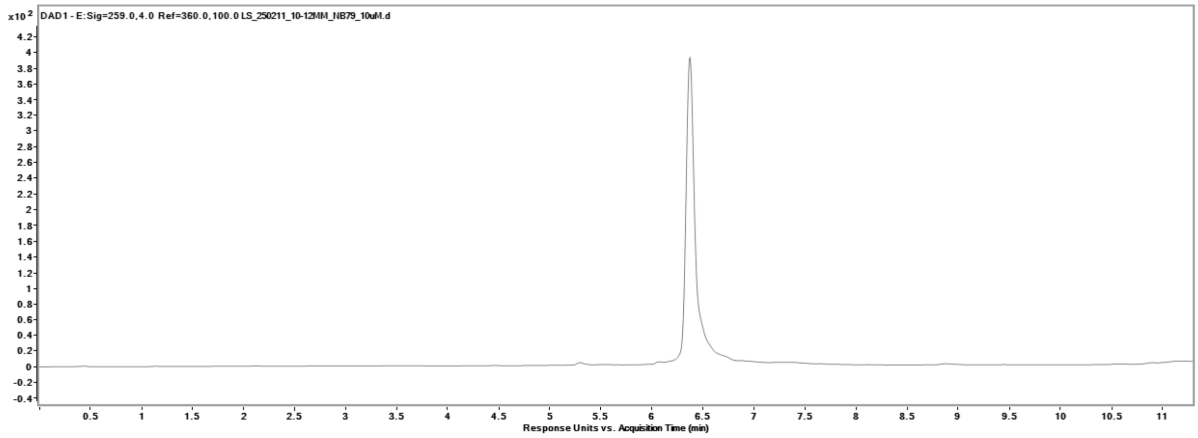

**GL-O TM13-15**

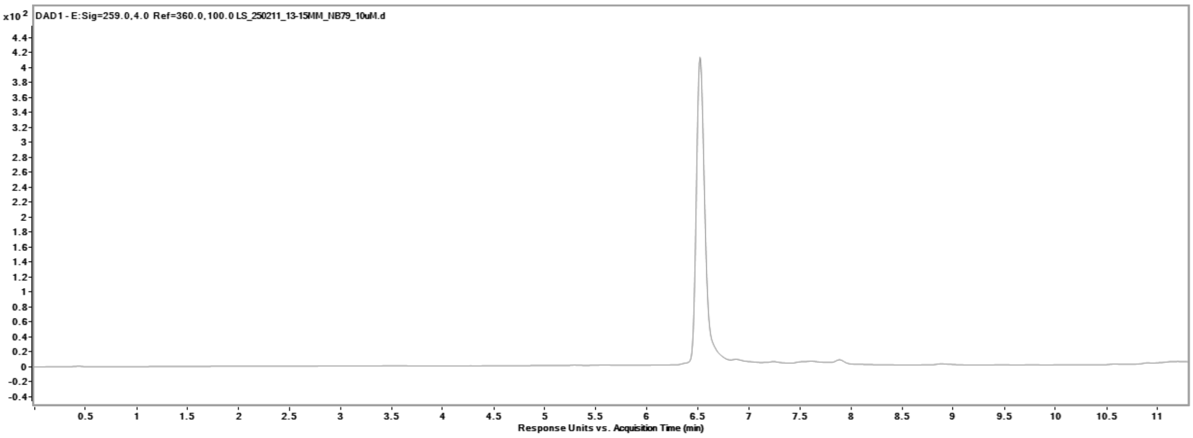

**Table S3.** Yields and mass of the GL-O conjugates.

| GL-O name | Yield | Calc. MS (g/mol) | Found MS (m/z) |
|---|---|---|---|
| **GL-O D6** | 66% | 2611,2446 | 2608,7248 |
| **GL-O D8** | 82% | 3217,6661 | 3215,8500 |
| **GL-O D10** | 57% | 3844,0856 | 3841,9774 |
| **GL-O D12** | 33% | 4452,4751 | 4452,0618 |
| **GL-O D14** | 43% | 5068,8957 | 5068,1141 |
| **GL-O D15** | 48% | 5398,1044 | 5397,2420 |
| **GL-O D16** | 30% | 5702,2992 | 5699,2611 |
| **GL-O D18** | 33% | 6344,7177 | 6344,3184 |
| **GL-O D20** | 54% | 6952,1232 | 6951,3930 |
| **GL-O P6** | 2% | 2289,0450 | 2290,0500 |
| **GL-O P8** | 6% | 2791,2788 | 2792,2542 |
| **GL-O P10** | 5% | 3341,4749 | 3342,4775 |
| **GL-O P12** | 5% | 3873.6779 | 3875,6694 |
| **GL-O P15** | 4% | 4691.0008 | 4692,9976 |
| **GL-O TM1-3** | 38% | 5389,0894 | 5373,1840 |
| **GL-O TM4-6** | 33% | 5373,0904 | 5390,1644 |
| **GL-O TM7-9** | 31% | 5391,0574 | 5388,2091 |
| **GL-O TM10-12** | 33% | 5361,1234 | 5386,1752 |
| **GL-O TM13-15** | 46% | 5365,0594 | 5364,1672 |

***Supplementary figures***

**Table S4.** Three site complementarity of a 13-nt oligonucleotide binding sequence of *C-Myc* Pu27.

| Chromosome | Matching site (gene) | Functions | Gene ID | Start-end position | Annotation | Sequence (flanking ±25)* |
|---|---|---|---|---|---|---|
| 8 | MYC proto-oncogene | - | ENSG00000136997 | 127735951 - 127735964 | Promoter | CGCCCTCTGCTTTGGGAACCCGG GAGGGGCGCTTATGGGGAGGGT GGGGAGGGTGGGGAAGGT |
| 1 | LMX1A (LIM homeobox transcription factor 1 alpha) | Transcription factor regulates gene expression involved in the development of neurons and pancreas. | ENSG00000162761 | 165213724 - 165213737 | Promoter | AAGGAACTGCTGAGGAAGGCAA GGACCATAAGCGCCCCAAACGTC CGAGAACCATCTTGACAA (antisense) TTGTCAAGATGGTTCTCGGACGT TTGGGGCGCTTATGGTCCTTGCC TTCCTCAGCAGTTCCTT |
| 12 | ABTB3 (ankyrin repeat and BTB domain containing 3) | Specifically expressed in cortical and hippocampal inhibitory interneurons and support in synaptic functions. | ENSG00000151136 | 107523249 - 107523262 | Intronic | AGTCAGTAGCTGATCCAAGCCAT AGGGGGCGCTTATGGCCACTAG AGAGCTACTTTCCTGGCA |
| 2 | CASC11 (LncRNA) | Cancer susceptibility candidate involved in gene regulation and found to be overexpressed in various cancers. | - | 17657528-17657585 | Intergenic | AAACGAAGTGAACTGTATCTGGG GCGCTTATGGAAAACAAACCCTC TAGCAGACTCAC |

*Sequence highlighted in red and G4-forming sequence is underlined.

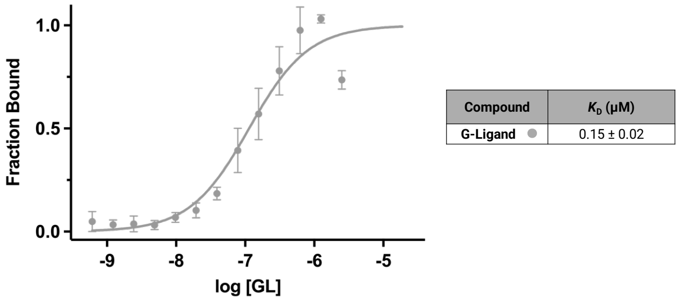

**Figure S1.** Binding affinity of the G4-ligand (GL). Dose-response curves obtained using MST where the GL was serial diluted to the G4 DNA 5´-labelled with a fluorescent tag. Dissociation constants ($K_D$) values and error bars correspond to two independent measurements.

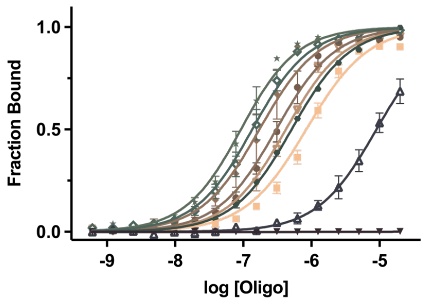

| Compound | $K_D$ (µM) |
|---|---|
| Varying length - DNA | |
| Oligo D6 ▼ | No binding |
| Oligo D8 △ | 9.16 ± 2.24 |
| Oligo D10 ● | 0.50 ± 0.01 |
| Oligo D12 ◇ | 0.13 ± 0.03 |
| Oligo D14 ★ | 0.08 ± 0.02 |
| Oligo D15 ● | 0.30 ± 0.06 |
| Oligo D16 ◆ | 0.17 ± 0.07 |
| Oligo D18 ▽ | 0.41 ± 0.05 |
| Oligo D20 ■ | 0.93 ± 0.12 |

**Figure S2.** Binding affinities of the oligonucleotides **D6**-**D20**. Dose-response curves obtained using MST where the oligonucleotides without G4-ligands (still contains a 5´hexylamine). **D6**-**D20** were serial diluted to the G4 DNA 5´-labelled with a fluorescent tag. Dissociation constants ($K_D$) values and error bars correspond to two independent measurements.

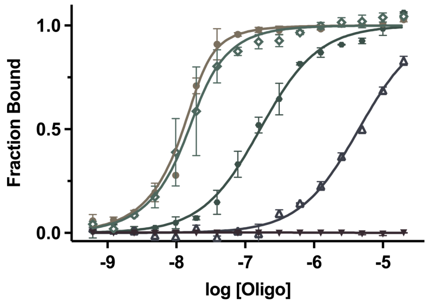

| Compound | $K_D$ (µM) |
|---|---|
| Varying length - PNA | |
| Oligo P6 ▼ | No binding |
| Oligo P8 △ | 4.46 ± 0.15 |
| Oligo P10 ● | 0.16 ± 0.04 |
| Oligo P12 ◇ | 7 ± 6 (nM) |
| Oligo P15 ● | 3 ± 2 (nM) |

**Figure S3.** Binding affinities of the oligonucleotides **P6**-**P15**. Dose-response curves obtained using MST where the oligonucleotides without G4-ligands (still contains a 5´hexylamine). **P6**-**P15** were serial diluted to the G4 DNA 5´-labelled with a fluorescent tag. Dissociation constants ($K_D$) values and error bars correspond to two independent measurements.

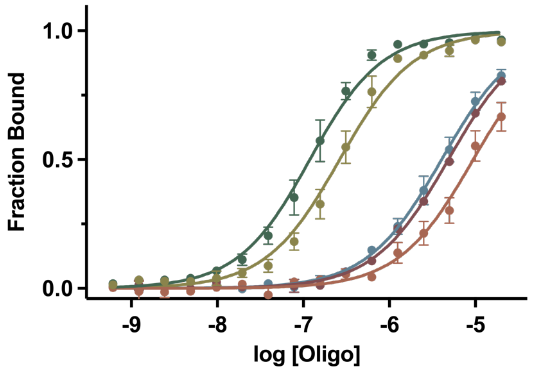

| Compound | $K_D$ (µM) |
|---|---|
| Varying triple mutations | |
| Oligo TM1-3 ● | 0.11 ± 0.03 |
| Oligo TM4-6 ● | 9.59 ± 2.34 |
| Oligo TM7-9 ● | 4.84 ± 0.17 |
| Oligo TM10-12 ● | 4.08 ± 0.74 |
| Oligo TM13-15 ● | 0.26 ± 0.05 |

**Figure S4.** Binding affinities of the oligonucleotides **TM1-3, 4-6, 7-9, 10-12, 13-15.** Dose-response curves obtained using MST where the oligonucleotides without G4-ligands (still contains a 5´hexylamine). **TM1-3, 4-6, 7-9, 10-12, 13-15** were serial diluted to the G4 DNA 5´-labelled with a fluorescent tag. Dissociation constants ($K_D$) values and error bars correspond to two independent measurements.

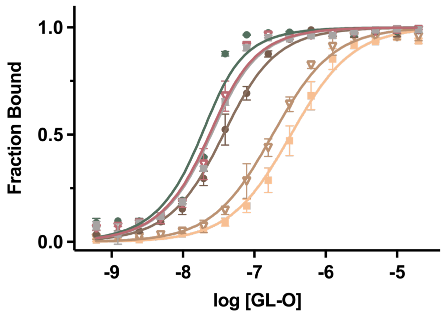

| Compound | $K_D$ (nM) |
|---|---|
| No incubation | |
| GL-O D15 ● | 27 ± 6 |
| GL-O D18 ▽ | 170 ± 34 |
| GL-O D20 ■ | 320 ± 80 |
| Incubation 24 h | |
| GL-O D15 ● | 8 ± 0 |
| GL-O D18 ▽ | 14 ± 4 |
| GL-O D20 ■ | 15 ± 1 |

**Figure S5.** Binding affinities of the GL-Os **D15**-**D20** with or without incubation in room temperature for 24 h. Dose-response curves obtained using MST. **D15**-**D20** were serial diluted to the G4 DNA 5´-labelled with a fluorescent tag. Dissociation constants ($K_D$) values and error bars correspond to two independent measurements.

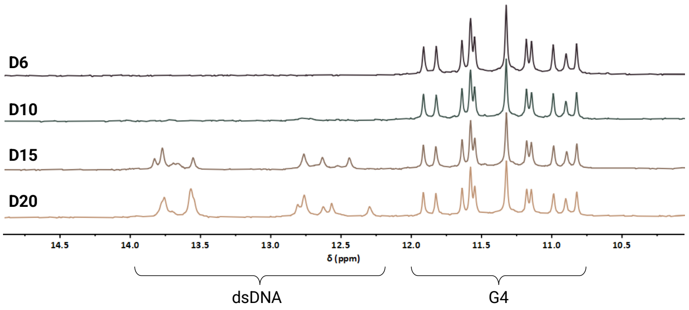

**Figure S6.** Binding interaction of the oligonucleotides **D6**-**D20** (still contains a 5´hexylamine) measure using $^1$H NMR. Spectra were recorded of *c-MYC* Pu24T with complementary flanking sequence in a 1:1 molar ratio at 25 °C. G4 imino signals appear between 10-12 ppm and double-stranded DNA signals appear between 12-14 ppm.

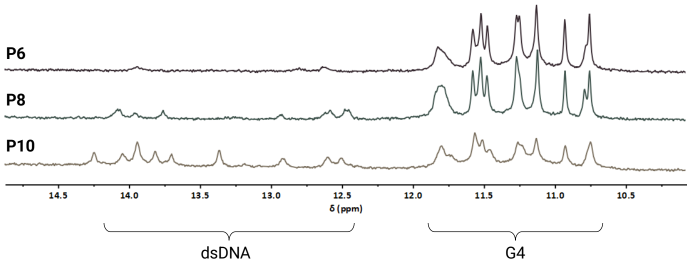

**Figure S7.** Binding interaction of the oligonucleotides **P6**-**P20** (still contains a 5´hexylamine) measure using $^1$H NMR. Spectra were recorded of *c-MYC* Pu24T with complementary flanking sequence in a 1:1 molar ratio at 25 °C. G4 imino signals appear between 10-12 ppm and double-stranded DNA signals appear between 12-14 ppm.

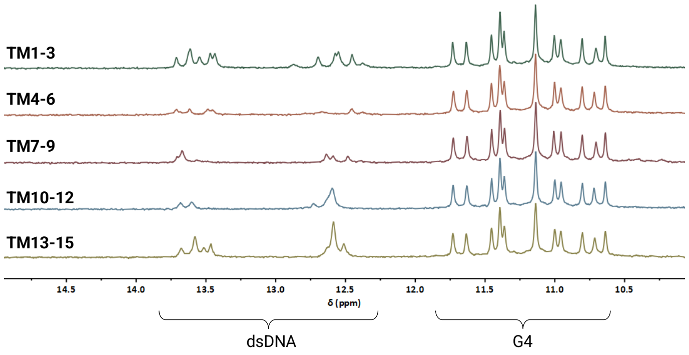

**Figure S8.** Binding interaction of the oligonucleotides **TM 1-3, 4-6, 7-9, 10-12, 13-15** (still contains a 5´hexylamine) measure using $^{1}$H NMR. Spectra were recorded of *c-MYC* Pu24T with complementary flanking sequence in a 1:1 molar ratio at 25 °C. G4 imino signals appear between 10-12 ppm and double-stranded DNA signals appear between 12-14 ppm.

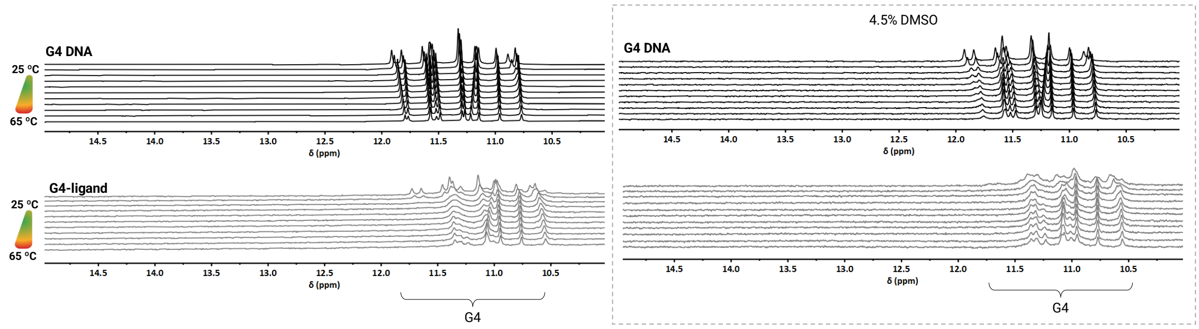

**Figure S9.** Thermal stabilization data of the G4 DNA alone and G4-ligand alone with and without DMSO using $^{1}$H NMR. Spectra were recorded of *c-MYC* Pu24T with complementary flanking at 25 and 35 °C and then with a temperature ramp of 2.5 °C from 45-65 °C. G4 imino signals appear between 10-12 ppm and double-stranded DNA signals appear between 12-14 ppm.

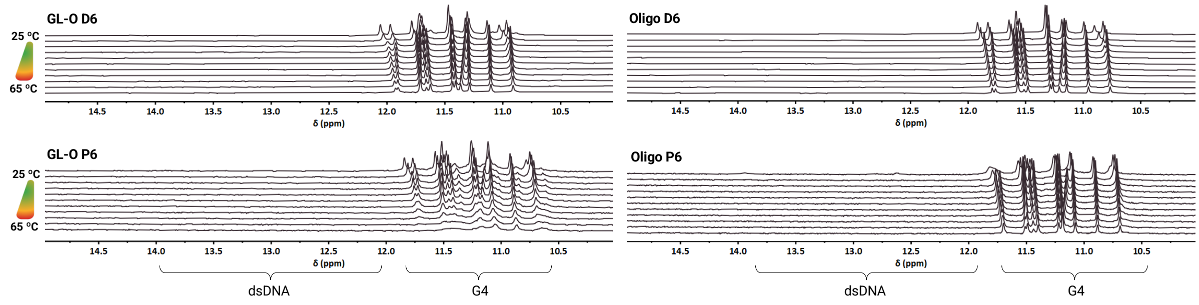

**Figure S10.** Thermal stabilization data of the GL-Os **D6** and **P6** with their unconjugated oligonucleotides using $^{1}$H NMR. Spectra were recorded of *c-MYC* Pu24T with complementary flanking at 25 and 35 °C and then with a temperature ramp of 2.5 °C from 45-65 °C. G4 imino signals appear between 10-12 ppm and double-stranded DNA signals appear between 12-14 ppm.

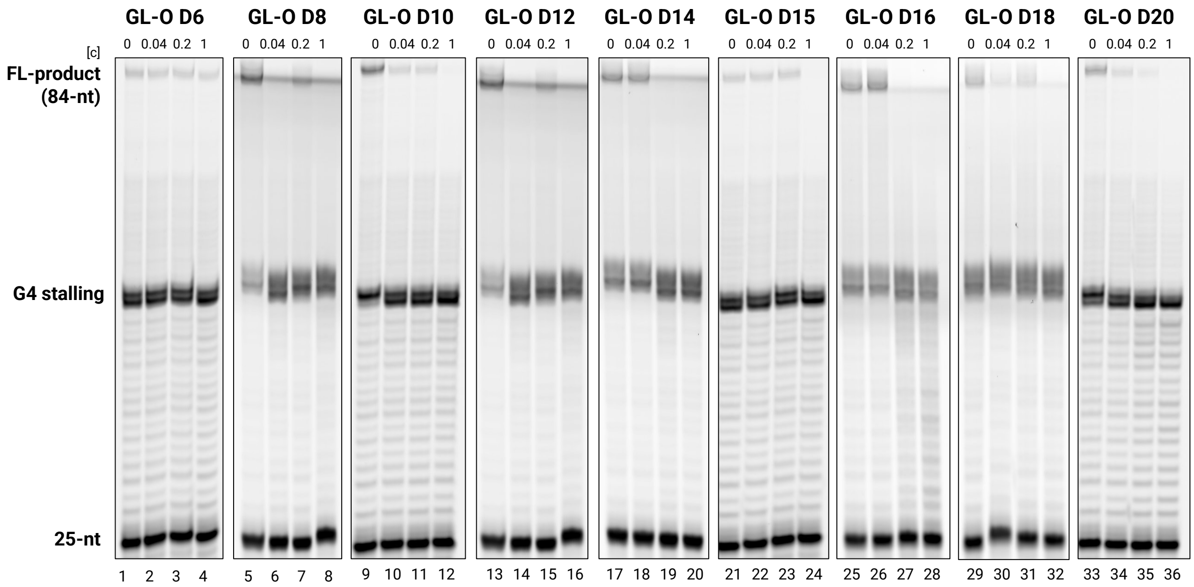

**Figure S11.** Taq Polymerase stabilization of the G4 DNA in presence of varying length oligonucleotide **GL-Os D6**-**D20**. Quantification of the Taq polymerase stop assay shown in as the full-length (FL, 84-nt) DNA product is expressed as a % of the full-length band intensity that was observed in the control reaction containing only the G4 template. Data represent the mean ± standard deviation from three independent experiments.

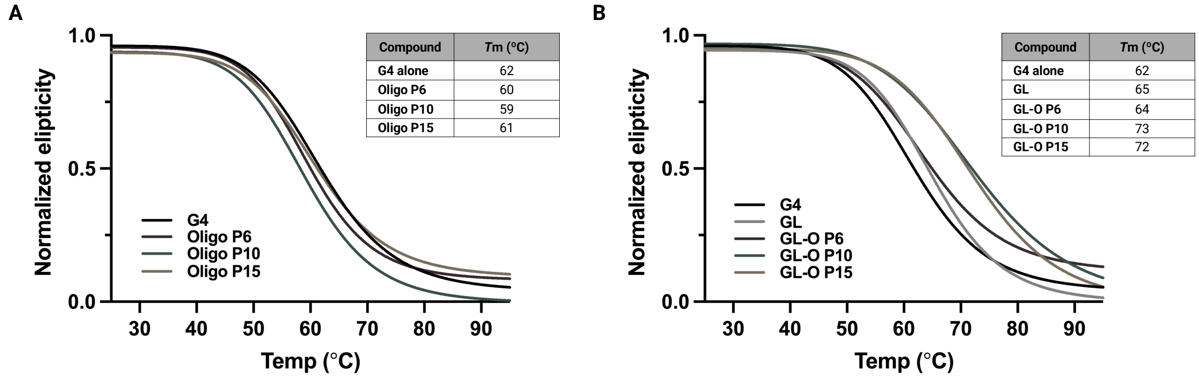

**Figure S12.** CD melting profiles of varying length oligonucleotide GL-Os **P6**-**P15** and unconjugated oligonucleotides **P6**-**P15.** Melting profiles were measured from 25-95 °C.

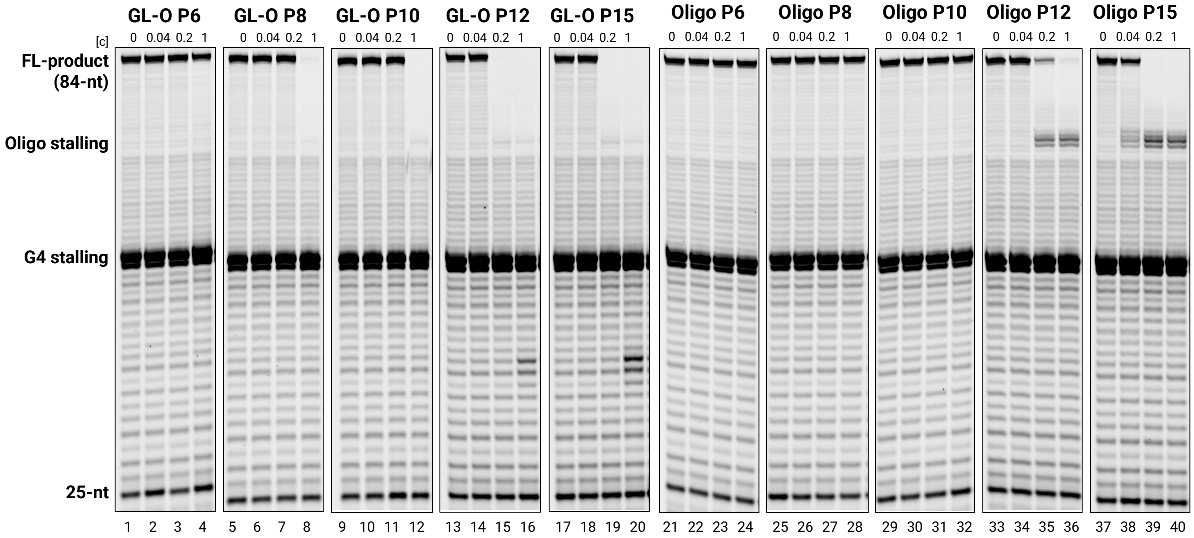

**Figure S13.** Taq Polymerase stabilization of the G4 DNA in presence of varying length oligonucleotide GL-Os **P6**-**P15** (lanes 1-20) and unconjugated oligonucleotides **P6**-**P15** (lanes 21-40). Quantification of the Taq polymerase stop assay shown in as the full-length (FL, 84-nt) DNA product is expressed as a % of the full-length band intensity that was observed in the control reaction containing only the G4 template. Data represent the mean ± standard deviation from three independent experiments.

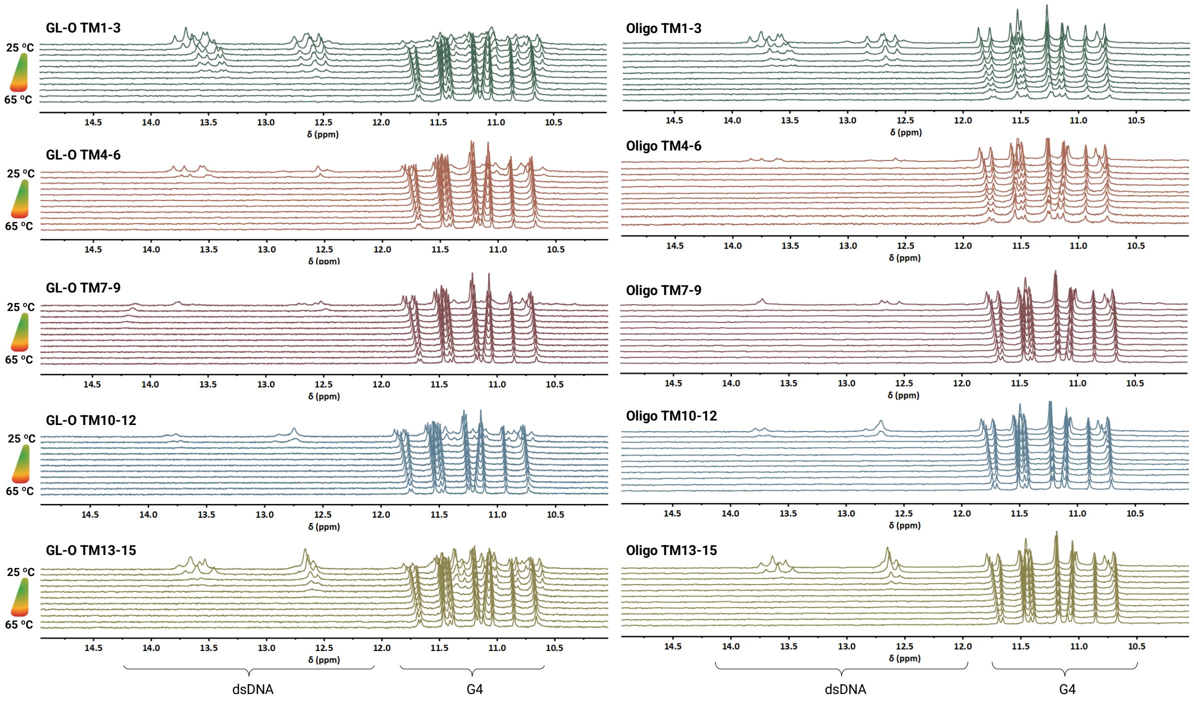

**Figure S14.** Thermal stabilization data of the GL-Os and unconjugated oligonucleotides **TM 1-3, TM4-6, TM7-9, TM10-12, TM13-15** using $^{1}$H NMR. Spectra were recorded of *c-MYC* Pu24T with complementary flanking at 25 and 35 °C and then with a temperature ramp of 2.5 °C from 45-65 °C. G4 imino signals appear between 10-12 ppm and double-stranded DNA signals appear between 12-14 ppm.

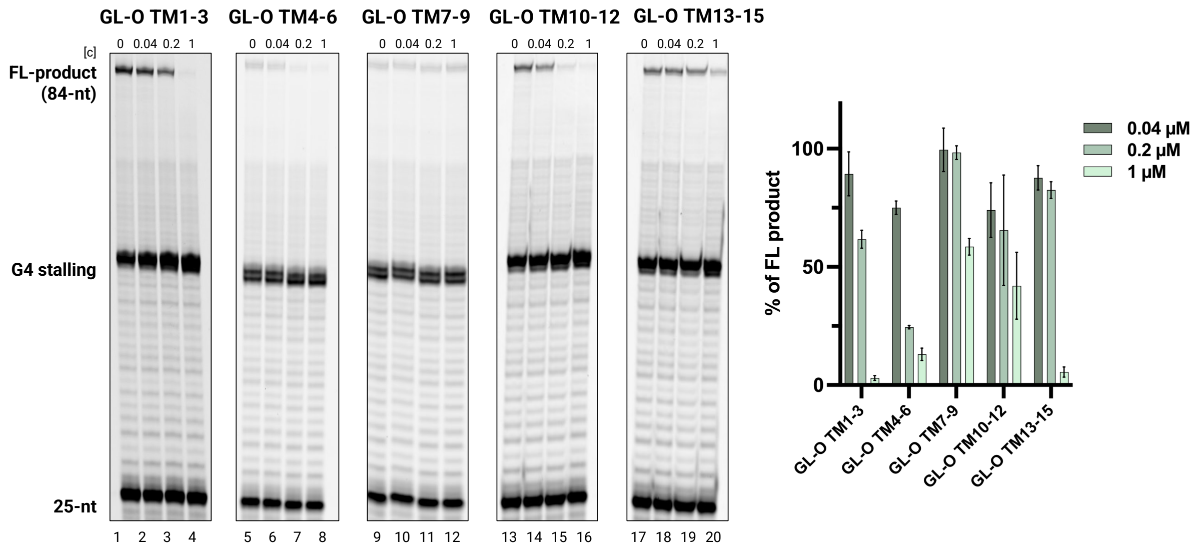

**Figure S15.** Taq Polymerase stabilization of the G4 DNA in presence of varying length oligonucleotide **GL-Os TM1-3**, **TM4-6**, **TM7-9**, **TM10-12** and **TM13-15**. Quantification of the Taq polymerase stop assay shown in as the full-length (FL, 84-nt) DNA product is expressed as a % of the full-length band intensity that was observed in the control reaction containing only the G4 template. Data represent the mean ± standard deviation from three independent experiments.

#### *DNA Oligonucleotide conjugations*

All oligonucleotides received through a collaboration with the Oligonucleotides and Targeted Delivery group, Discovery Sciences at AstraZeneca R&D (Gothenburg, Sweden) or purchased from IDT (Integrated DNA Technologies, Sweden). 5´-hexylamine oligonucleotide ($Na^+$ salt, 50 nmol) was dissolved in 0.1 M $NaHCO_3$ (aq) (50 µL, 1 mM). A solution of (1R,8S,9s)-bicyclo[6.1.0]non-4-yn-9-ylmethyl N-succinimidyl carbonate (BCN-NHS ester) (20 equiv., 5 µL, 0.2 M) was freshly prepared and added to the oligonucleotide solution. The reaction solution was vortexed and shaken for 24 h. Followed by precipitation of oligonucleotide conjugate using 10% v/v of 3 M NaOAc and 4 times the volume EtOH and kept in freezer for 30 min. Then the solution was centrifuged for 15 min followed by removing the supernatant. The precipitate pellet was resuspended in the same volume of NaOAc and EtOH and repeated the precipitation. After this, the pellet was washed with only EtOH followed by drying the pellet by gently adding $N_2$ gas and then resuspended in 0.1 M $NaHCO_3$ (aq) (50 µL, 1 mM). The G4-ligand baring an azide functionality (previously described)[1] (2.5 equiv., 25 µL, 10 mM in DMSO), was added to the dissolved oligonucleotide and vortexed and shaken for 24 h. The reaction mixture was dissolved in more water and DMSO and filtered before purified by RP-HPLC. Purification was done using a C18 semi-preparative column and 2 mL/min flowrate and a linear gradient of 5 to 60% solvent B (100% ACN). Solvent A was a 50 mM triethylammonium acetate buffer. The conjugation products were confirmed with HRMS ($ESI^-$).

#### *Peptide Nucleic Acid synthesis and conjugations*

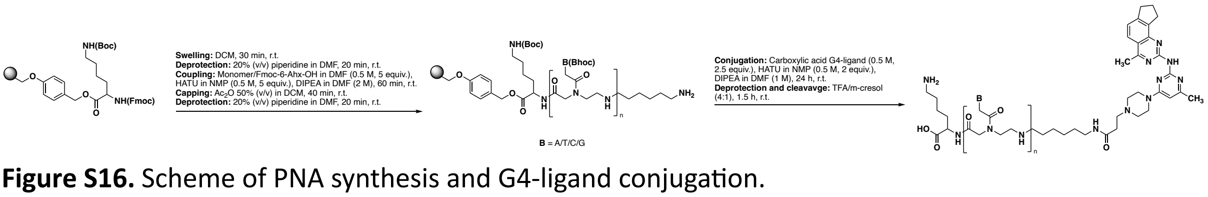

**Figure S16.** Scheme of PNA synthesis and G4-ligand conjugation.

All reagents and solvents including resins, piperidine, hexafluorophosphate azabenzotriazole tetramethyl uronium (HATU), *N*,*N*-Diisopropylethylamine (DIPEA), trifluoro acetic acid (TFA), m-cresol, *N*-Methyl-2-pyrrolidone (NMP), acetonitrile (ACN), formic acid, diethyl ether was purchased from *Merck Sigma Aldrich*. The Fmoc-PNA(Bhoc)-OH monomers were purchased from *Iris Biotech GmbH*, Fmoc-6-Ahx-OH was purchased from *Chemtronica,* Peptide Grade dimethylformamide (DMF) was purchased from *CARLO ERBA Reagents*. The PP-reactors were purchased from *Biotage*. RP-HPLC was carried out on a Hitachi HPLC system using a Phenomenex Luna® Omega 5 µm Polar C18 100Å (250x 10 mm) column.

Peptide grade DMF was used for PNA synthesis. The reported equivalents are relative to the amount of resin, and all reactions were performed at room temperature.

The PNAs were synthesized on a 5-20 µmol scale on a Syro I automated peptide synthesizer (Biotage) using preloaded resin Fmoc-Lys(Boc)-Wang resin (loading 0.61 mmol/g) for **P6-P10** and Fmoc-Lys(Boc)-Wang LL (loading 0.2 – 0.4 mmol/g) for **P12-P15**. The PNA synthesis was performed in PP reactors with PE frit, and deprotection and cleavage were performed in PP reactors with PTFE frit.

The resin was allowed to swell in DCM for 30 min under agitation. Fmoc deprotection cycle was performed with 20% (v/v) piperidine in DMF for 15 min. The Fmoc-PNA(Bhoc)-OH monomers (0.5 M in DMF, 5 eq) were coupled for 1 h using HATU (0.5M in NMP, 5 eq), DIPEA (2.0 M in DMF) for 60 minutes. A capping step was performed after each coupling using $Ac_2O$ 50% (v/v) in DCM. After PNA elongation, the N-terminal was coupled with Fmoc-6-Ahx-OH, followed by Fmoc deprotection according to the aforementioned procedures. Conjugation to the G4-ligand was performed on a 5 µmol scale. The N-terminal amine of the monomers was coupled with G4-ligand (0.25 M in NMP, 2.5 eq), HATU (0.25 M in NMP, 5 eq), DIPEA (1.0 M in DMF), and the coupling time was extended to 24 h.

After the synthesis was completed, the resin was suspended in a mixture of TFA:m-cresol (4:1) and vortexed for 1.5 h, resulting in simultaneous deprotection and cleavage from the resin. The solution was filtered into ice-cold $Et_2O$ and centrifuged. The white precipitate was purified using RP-HPLC. Purification was carried out at room temperature using detection at 260 nm on a C18 semi-preparative column with a 2.5 mL/min flow rate in a gradient system of 0-20% B over 24 min then 20-100% B over 2 min (B 100% ACN and (A 0.1 Formic acid in Milli-Q water) for **P6-P10** and gradient system of 0-30% over 11 min B then 30-100% B over 15 min (B 0.1% TFA in ACN and A 0.1% TFA in Milli-Q water). The conjugated products were confirmed with HRMS ($ESI^+$).

**Table S5.** Protocols for the PNA synthesis.

| Step | Reagent/solvent | Volume (µL) | Cycles x Time |
|---|---|---|---|
| Swelling | DCM | 500 | 30 minutes |
| Wash | DMF | 5 x 500 | 5 x 1 minute |
| Deprotection | 20% (v/v) piperidine in DMF | 500 | 2 x 3 minutes<br>1 x 9 minutes |
| Wash | DMF | 5 x 500 | 5 x 1 minute |
| Coupling | Monomer in DMF<br>HATU in NMP<br>DIPEA in DMF | 1 x 200<br>1 x 210<br>1 x 100 | 60 minutes |
| Wash | DMF | 5 x 500 | 5 x 1 minute |
| Capping | Ac2O 50% (v/v) in DCM<br>DIPEA | 1 x 200<br>1 x 100 | 40 minutes |
| Wash | DMF | 5 x 500 | 5 x 1 minute |
| Conjugation of G4-Ligand | G4-Ligand in NMP<br>HATU in NMP<br>DIPEA in DMF | 1 x 200<br>1 x 210<br>1 x 100 | 24 h |

***In vitro* assays**

**Microscale thermophoresis (MST).** The G4 template consisting of the G4-forming sequence and the flanking sequence was purchased with a Cy5 5´-label. The G4 DNA was annealed in MST buffer (10 mM potassium phosphate, 100 mM KCl, 0.05% Tween 20, pH 7.4) by heating at 96 °C for 5 min followed by cooling down to room temperature before storing in the fridge overnight. All MST experiments were done on a Monolith NT.115 (Nanotemper, Germany) instrument and performed in MST buffer with standard Monolith capillaries. The G4 DNA concentration was kept constant at 20 nM, and the GL-Os were serial diluted (1:1) with highest concentration of 5-40 µM. For PNA-based GL-Os, binding experiments were conducted both in the presence and absence of DMSO, and no differences in thermophoresis traces or derived binding affinities were observed. MST traces and binding affinity constants ($K_D$) were obtained using the Monolith analysis software and plotted and visualized in GraphPad Prism 10.

**Proton nuclear magnetic resonance spectroscopy ($^1$H NMR)**. G4 DNA was prepared by annealing in 10 mM potassium phosphate buffer (3 mM KCl, 110 µM G4 DNA, pH 7.4). The solution was heated to 96 °C for 5 minutes, then gradually cooled to room temperature and stored overnight at 4 °C. Prior to NMR analysis, 10% $D_2O$ was added to the G4 DNA solution to achieve a final concentration of 100 µM, and the sample was transferred to a 3 mm NMR tube. GL-O (1 mM) was added to the DNA solution in two steps: first, 2 µL (0.5 equivalents) was added, and after 10 minutes, a $^1$H NMR spectrum was recorded. A second aliquot (0.5 equivalents) was then added to achieve a 1:1 molar ratio with G4 DNA, and another spectrum was recorded. For DNA-based GL-Os, samples were prepared in aqueous buffer, whereas PNA-based constructs required the addition of DMSO to a final concentration of 4.5% (v/v) to ensure complete solubility. NMR experiments were conducted on an 850 MHz Bruker AVANCE III HD spectrometer equipped with a 5 mm TCI cryoprobe at 298 K. The transmitter frequency offset (O1P) was set at 4.7 ppm, with a spectral width of 22 ppm. One-dimensional $^1$H NMR spectra were acquired using excitation sculpting with 512 scans per spectrum. For thermal stability studies, samples were equilibrated at temperatures ranging from 313 K to 338 K in 2.5–10 K intervals. Spectra were recorded after a 5-minute equilibration at each temperature. Data were processed using MestreNova version 10.0.2.

NMR experiments of PNA GL-Os **P6**-**P15** were done in 4.5% DMSO by adding DMSO-$d_6$ (20 µL) and $D_2O$ (10 µL) to a G4 DNA solution (170 µL), to achieve a final concentration of 100 µM G4 DNA.

**Taq polymerase STOP assay.** Taq Polymerase stop assay was adapted from Berner et al.[2] DNA templates were annealed to fluorescent labelled primers in 100 mM KCl by heating to 95 °C for 5 minutes followed by slow cooling to room temperature. The indicated compound concentrations were added to 40 nM annealed template in 1x Taq Buffer (10 mM Tris-HCl pH 8.8, 50 mM KCl, Thermo Fisher Scientific), 1.5 mM $MgCl_2$, and 0.05 U/µL Taq Polymerase (Thermo Fisher Scientific). For PNA-based GL-Os, assays were performed both in the presence and absence of DMSO, and no differences in polymerase stalling or full-length product formation were observed. Samples were preincubated on ice (10 minutes) and reactions initiated with the addition of dNTPS (100 µM) and transferring the samples to 37 °C. After 15 min at 37 °C, reactions were stopped by addition of equal volume of 2x stop solution (0.5% SDS, 25 mM EDTA, XC-Dye in Formamide) and separated on a 12% polyacrylamide Tris-Borate-EDTA (TBE) gel containing 25% formamide and 8 M urea. Fluorescent signal was detected with a Typhoon Scanner (Amersham Biosciences). The intensity of the full-length band was quantified using Image Quant TL 10.2 software (GE Healthcare Life Sciences) and compared to sample without compound.

The PNA GL-Os **P6**-**P15** were run in a final concentration of 0.45% DMSO in the assay.

***In vitro* endonuclease digestion.** Stability of oligos against Mung bean nuclease digestion was assessed. 4 µM oligos were incubated with 3 units of enzyme (MBN, NEB) in 1X MBN buffer (MBN) at 30 degrees. At indicated time points, reactions were quenched by adding DNA Gel loading dye (50 mM EDTA, 50 mM HEPES, 30% Glycerol, 0.001% Bromophenol Blue) and heating at 95 degrees. Digestion products were analysed by polyacrylamide gel electrophoresis (PAGE). DNA samples were resolved on 16% TBE gels for 1 h at 100 V in 1X TBE buffer. PNA samples were loaded on 4-20% SDS PAGE gels (Mini-PROTEAN TGX Gels, Bio-Rad) and run for 1 h at 120 V in 1X SDS running buffer (Tris Glycine). TBE gels were stained with Gel red (Biotium), while SDS PAGE gels were stained with Instant Blue (Abcam). Gels were imaged using a ChemiDoc Imaging system (Bio-Rad).

**CD melting temperature assay.** G4 (3 µM) was folded in K-phosphate buffer (10 mM, pH 7.4) containing KCl (3 mM) by heating for 5 minutes at 95°C, then allowed to cool to room temperature. PNA in DMSO was added in a 1:1 ratio, resulting in a final DMSO concentration of 0.3%. CD measurements were performed on a J-1700 Circular Dichroism Spectrophotometer (Jasco International Co. Ltd.). A quartz cuvette with a path length of 1 mm was used for measurements. CD spectra were recorded between λ = 200 – 350 nm with an interval of 0.5 nm and a scan rate of 100 nm/min. Thermal melting was recorded between 25°C and 95°C with increments of 10°C at a speed of 1 °C/min

***Computational assessment***

**Sequence alignment analysis.** The specificity of 5′ G4 flanking sequences of varying lengths was assessed by aligning these DNA sequences to the human reference genome. The human reference genome (GRCh38) was retrieved from the UCSC Genome Browser,[3] and flanking sequence was obtained from NCBI nucleotide database. FASTA datasets containing flanking sequences of 5–20 bases were generated and each dataset was independently aligned to the human reference genome using Bowtie 2.0,[4] with no mismatch allowed. The aligned reads in SAM files were converted to BAM format and subsequently sorted and indexed using SAMtools[5] with default settings. The final BAM files were used for downstream sequence annotation and comparative analysis of mapping specificity across different flanking lengths.

***References***

1. Bhuma, N., Chand, K., Andréasson, M., Mason, J., Das, R.N., Patel, A.K., Öhlund, D., and Chorell, E. (2023). The effect of side chain variations on quinazoline-pyrimidine G-quadruplex DNA ligands. Eur. J. Med. Chem. *248*, 115103.
2. Berner, A., Das, R.N., Bhuma, N., Golebiewska, J., Abrahamsson, A., Andréasson, M., Chaudhari, N., Doimo, M., Bose, P.P., Chand, K., et al. (2024). G4-Ligand-Conjugated Oligonucleotides Mediate Selective Binding and Stabilization of Individual G4 DNA Structures. J. Am. Chem. Soc. *146*, 6926-6935.
3. Casper, J., Speir, M.L., Raney, B.J., Perez, G., Nassar, L.R., Lee, C.M., Hinrichs, A.S., Gonzalez, J.N., Fischer, C., Diekhans, M., et al. (2025). The UCSC Genome Browser database: 2026 update. Nucleic Acids Research *54*, D1331-D1335.
4. Langmead, B., and Salzberg, S.L. (2012). Fast gapped-read alignment with Bowtie 2. Nature Methods *9*, 357-359.
5. Li, H., Handsaker, B., Wysoker, A., Fennell, T., Ruan, J., Homer, N., Marth, G., Abecasis, G., Durbin, R., and Subgroup, G.P.D.P. (2009). The Sequence Alignment/Map format and SAMtools. Bioinformatics *25*, 2078-2079.
